# Discovery, functionality and global dynamics of *Bifidobacterium longum* subsp. *nexti*: a novel starch-degrading subspecies in the infant gut

**DOI:** 10.1101/2025.06.23.661074

**Authors:** Dena Ennis, Sergio Andreu-Sánchez, Yuvashankar Kavanal Jayaprakash, Trishla Sinha, Haoran Peng, Omer Goldberger, James A. D. Docherty, Nataliia Kuzub, Asier Fernández-Pato, Shijie Zhao, Yan Shao, Yarden Levin, LLNEXT cohort study, Orna Amster-Choder, Trevor D. Lawley, Rinse K. Weersma, Eric Alm, Maria Carmen Collado, Jingyuan Fu, Alexander Kurilshikov, Hermie J.M. Harmsen, Cyrus A. Mallon, Douwe van Sinderen, Alexandra Zhernakova, Moran Yassour

**Affiliations:** Microbiology & Molecular Genetics Department, Faculty of Medicine, The Hebrew University of Jerusalem, Jerusalem, Israel; Department of Genetics, University of Groningen and University Medical Center Groningen, The Netherlands; Department of Microbiology, University of Groningen and University Medical Center Groningen, The Netherlands; Department of Biological Engineering, Massachusetts Institute of Technology, Cambridge, MA, USA; Center for Microbiome Informatics and Therapeutics, Massachusetts Institute of Technology, Cambridge, MA, USA; Host–Microbiota Interactions Laboratory, Wellcome Sanger Institute, Hinxton, UK; See supplementary information for authors and their affiliations; Department of Gastroenterology and Hepatology, University of Groningen and University Medical Center Groningen, The Netherlands; The Broad Institute of MIT and Harvard, Cambridge, MA, USA; Department of Biotechnology, Institute of Agrochemistry and Food Technology-National Research Council, Valencia, Spain; APC Microbiome Institute and School of Microbiology, University College Cork, Cork T12 YT20, Ireland; The Rachel and Selim Benin School of Computer Science and Engineering, The Hebrew University of Jerusalem, Jerusalem, Israel; The Hebrew University Computational Medicine Center, Faculty of Medicine, The Hebrew University of Jerusalem, Jerusalem, Israel

## Abstract

*Bifidobacterium longum* is a foundational, genetically diverse member of the infant gut microbiome, yet resolving its subspecies in metagenomic data remains challenging. Here, we performed a deep phylogenetic characterization of the *B. longum* species utilizing 4,577 fecal metagenomes from the LLNEXT mother-infant cohort alongside a global dataset of 1,220 genomes and metagenome-assembled genomes (MAGs). We identified a novel, geographically widespread lineage: *Bifidobacterium longum* subsp. *nexti*. Using a polyphasic taxonomic analysis, including average nucleotide identity and digital DNA-DNA hybridization, confirms subspecies-level divergence. Distinct from other members of the *B. longum* species, *BL. nexti* harbors α-amylopullulanase and grows on soluble starch and often pullulan, consistent with α-glucan utilization. Within the LLNEXT cohort, *BL. nexti* is enriched in formula-fed infants and shows evidence consistent with vertical transmission. Globally, while *BL. nexti* is geographically widespread, it usually persists at low relative abundance across diverse populations. By resolving a major fraction of previously unclassified *B. longum* diversity, this study provides new insights into the specialized lineages and ecological dynamics driving the assembly of the developing infant microbiome.

## Introduction

The genus *Bifidobacterium* is a cornerstone of the human gut microbiome, initiating colonization in early infancy and persisting as a prevalent inhabitant throughout the human lifespan^1,2^. This ubiquitous presence underscores a highly successful symbiotic relationship, where *Bifidobacterium* species contribute to host health by maintaining intestinal barrier integrity^3^, modulating immune system development^4^, and protecting against pathogen-induced colonic cell death^5^. In full-term, breastfed infants, *Bifidobacterium* species typically dominate the gut microbiome^6–8^. Conversely, their absence has been linked to various dysbiotic conditions, leading to their widespread administration as probiotics in vulnerable populations, such as preterm infants, to promote healthy microbial succession^9,10^.

Among the species inhabiting the human gut, *Bifidobacterium longum* is particularly significant since it is known as one of the most abundant species across all life stages^11–13^. The current taxonomic framework partitions *B. longum* into several subspecies that reflect specialization for distinct ecological niches^14,15^, including *B. longum* subsp. *longum* (*BL. longum*), *B. longum* subsp. *infantis* (*BL. infantis*), and *B. longum* subsp. *suis* (*BL. suis*)^16^. Although formally classified as separate subspecies, *B. longum* subsp. *suillum*^17^ and *B. longum* subsp. *iuvenis*^18^ exhibit high genomic and metabolic similarity to *BL. suis*, justifying their consolidation into a broader *BL. suis* clade^17,18^. Within this group, *BL. suillum* is characterized primarily as a urease-negative variant of the swine-associated *BL. suis*^17^, whereas *BL. iuvenis* represents a closely related lineage frequently isolated from the human gut^18,19^.

A defining feature of the *B. longum* species is its extensive intraspecies genomic diversity and metabolic specialization, driven primarily by differential carbohydrate utilization. Historically classified under a single species umbrella, a recent genomic atlas has proposed separating *BL. infantis* and *BL. longum* into two distinct species, with the latter further partitioned into specialized subspecies^20^. *BL. infantis* represents a hallmark of adaptation to the neonatal environment; it possesses a highly conserved ∼40 kb gene cluster specifically dedicated to the efficient transport and degradation of human milk oligosaccharides (HMOs)^21,22^. In contrast, *BL. longum* is generally adapted to the post-weaning and adult gut, exhibiting a metabolic repertoire specialized for plant-derived carbohydrates such as arabinoxylan and xylose^15,23^. While *BL. suis* was typically associated with pigs, it has recently been identified in the infant gut and shares plant-sugar utilization traits with *BL. longum*, alongside evidence of HMO utilization capabilities^18,19^. These observations indicate a genomic continuum between *BL. longum* and *BL. suis*, suggesting that their taxonomic distinctions are not absolute. The identification of *BL. longum* strains with mixed metabolic profiles, such as the ability to utilize fucosylated HMOs^24^, highlights significant genomic heterogeneity and suggests that ecological transitions and niche adaptations between these subspecies are complex^25^.

Accurately characterizing and quantifying fine-scale diversity within *B. longum* presents considerable challenges for metagenomic-based studies. Traditional taxonomic classifiers often struggle with the high degree of sequence conservation between subspecies^26^. For instance, methods using whole genomes as a reference database, such as k-mer based Kraken^27^, lead to high false-positive detection rates due to the unspecific subspecies genomic content^28^. In contrast, MetaPhlAn^29^, which relies on a set of clade-specific marker genes, currently treats *B. longum* as a single species-level genomic bin (SGB). Since it uses a single set of marker genes for the entire species, standard MetaPhlAn 4.0, which is the updated version at the moment of writing this paper, cannot distinguish between individual subspecies. To address this limitation, we previously developed a MetaPhlAn-based method by incorporating specific subspecies marker genes for *BL. infantis* and *BL. longum* into the reference database^30^. This enhancement allows for accurate differentiation and quantification of these two subspecies within complex metagenomic samples and can be further expanded to other subspecies.

Here, to improve our understanding of the diversity within *Bifidobacterium longum* and the ecological factors shaping its colonization of the infant gut, we performed a comprehensive phylogenomic analysis of 4,577 fecal metagenomes from the longitudinal LLNEXT mother-infant cohort. This analysis uncovered *Bifidobacterium longum* subsp. *nexti*, a previously undescribed *B. longum* subspecies, expanding the known diversity within this species. To enable its accurate detection in metagenomic datasets, we identified subspecies-specific marker genes that distinguish *BL. nexti* from other *B. longum* subspecies. Leveraging this enhanced taxonomic resolution, we characterized the global prevalence and colonization dynamics of *BL. nexti* alongside other *B. longum* subspecies across 6,162 samples from 18 geographically diverse cohorts. Finally, we explore its genomic content and metabolic potential to elucidate its functional role and potential survival strategies within the human gut environment.

## Results

### Phylogenetic and genomic analyses of *Bifidobacterium longum* **reveal a novel subspecies**

In this study we aimed to characterize the subspecies composition and diversity of *Bifidobacterium longum*, a keystone early-life microbe, within LLNEXT, a large Dutch longitudinal mother-infant cohort^31^. We profiled 4,577 fecal metagenomes from 714 mother-infant dyads, spanning early pregnancy through the first year of life. Using strain-level phylogenetic analysis together with relative abundance quantification (**Methods**), we identified four distinct *B. longum* lineages (**Figure 1A**). The largest clade contained the reference genomes of *BL. longum* and was dominated by samples with high *B. longum* subsp. *longum* (*BL. longum)* abundance. A second clade contained *B. longum* subsp. *infantis* (*BL. infantis)* reference genomes and correspondingly high *BL. infantis* abundance, supporting the accuracy of our subspecies assignments. A third clade comprised genomes previously annotated as *B. longum* subsp. *suis* (*BL. suis*; including *B. longum* subsp. *suillum* and *B. longum* subsp. *iuvenis*) and was characterized by elevated levels of unclassified *B. longum*, consistent with the close genomic similarity between *BL. longum* and *BL. suis*, which complicates their separation^32^. The fourth clade consisted predominantly of samples previously classified as unclassified *B. longum* and included nine reference genomes (**Figure 1A**, orange stars). Based on its distinct phylogenetic position, we propose this lineage as a novel subspecies, *Bifidobacterium longum* subsp. *nexti* subsp. nov. (*BL. nexti*), named after its discovery and isolation from an infant in the LLNEXT cohort.

**Figure 1:**
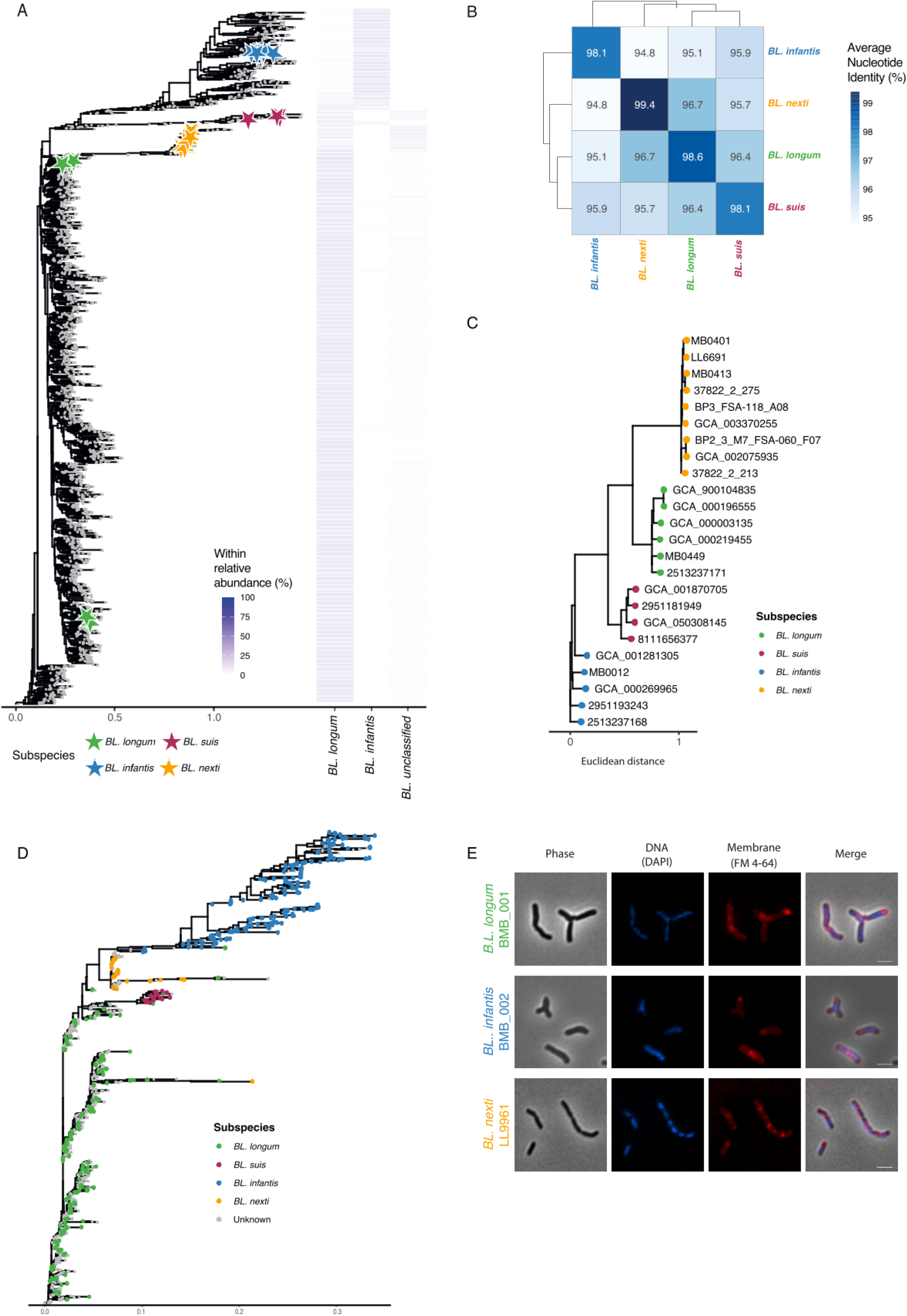
Phylogenomic analyses reveal four distinct *Bifidobacterium longum* lineages, including the novel subspecies *BL. nexti*. **(A)** Marker-gene-based phylogenetic tree (StrainPhlAn 4) of dominant gut strains in samples from the LLNEXT cohort. Stars represent reference genomes and are colored based on public subspecies annotation: *BL. longum*, *BL. infantis*, *BL. suis*, and the newly proposed *BL. nexti*. Within-species relative abundances are displayed as a heatmap on the right, showing abundances for subspecies *BL. longum*, *BL. infantis*, and unclassified *B. longum*. **(B)** Average genomic ANI (average nucleotide identity) between genomes and MAGs belonging to the different *B. longum* subspecies. Clusters support the presence of four distinct subspecies. Genomes are colored based on prior public subspecies annotations, while MAGs from the LLNEXT cohort are colored according to their sample annotation as defined in the phylogenetic tree in Figure 1A. **(C)** Dendrogram of selected genomes generated from digital DNA-DNA hybridization (dDDH) distances among 24 selected *B. longum* genomes. **(D)** Phylogenetic tree of all genomes and MAGs based on core genome SNVs (671 genes). Genomes and MAGs are colored as in (B). **(E)** Representative microscopic images of *BL. infantis*, *BL. longum*, and *BL. nexti* strains during the logarithmic growth phase. Cells are stained to visualize the cell membrane and DNA. Scale bar, 2.5 μm.

We next aimed to further characterize *BL. nexti* as a novel subspecies, by examining variation across 501 isolate genomes and 698 metagenome-assembled genomes (MAGs) of *B. longum* (**Supplementary Table 1**; **Methods**). Using average nucleotide identity (ANI)-based clustering we recovered the same four-lineage structure observed in the phylogenomic tree, with three clusters matching known *B. longum* subspecies and the fourth corresponding to *BL. nexti* (**Figure 1B; Supplementary Figure 1A; Supplementary Table 2**). The *BL. nexti* clade comprised nine genomes: seven that we isolated from human fecal samples, including five from infants in the Netherlands, Ireland, and the United Kingdom, and two from adults in the Netherlands, while the remaining two publicly available genomes originated from human milk^33^ and vaginal microbiomes^34^ in Italy and Canada, respectively (**Supplementary Table 3**). Mean ANI values between *BL. nexti* and the other subspecies ranged from 94.8% to 96.7%, below the commonly used 97% threshold for subspecies delineation^35^, while largely remaining within the range expected for members of the same bacterial species. In agreement with ANI, digital DNA-DNA hybridization (dDDH; **Methods**) values between *BL. nexti* and other *B. longum* subspecies ranged from 62% to 75.9%, below the suggested 79-80% subspecies boundary, whereas within-subspecies comparisons reached 94-99.3% (**Figure 1C; Supplementary Figure 1B; Supplementary Table 4**). Interestingly, *BL. infantis* exhibited low dDDH values compared to other subspecies (62-69.4%), supporting the notion that it is more distinct than other *B. longum* subspecies and may represent a different species, as has previously been suggested^20,21,36^. Both ANI and dDDH results indicated that *BL. nexti* is genomically closest to *BL. longum* and most divergent from *BL. infantis* (**Figure 1B,C**).

Phylogenetic reconstruction based on core-genome single nucleotide variants (SNVs) further confirmed the coherence of this proposed subspecies: all *BL. nexti* genomes and MAGs formed a single monophyletic clade, indicating recent common ancestry and clear separation from the other *B. longum* subspecies (**Figure 1D**). By contrast, analysis of complete 16S rRNA gene sequences using *B. breve* as an outgroup, confirmed that all selected genomes belong to the *B. longum* species complex (**Supplementary Figure 1C**). However, the 16S rRNA gene lacked sufficient resolution to distinguish between the different *B. longum* subspecies (**Supplementary Figure 1C**). Thus, while 16S rRNA supported species-level assignment, whole-genome approaches are required for subspecies-level discrimination.

Finally, phenotypic features further supported the distinctiveness of *BL. nexti*. We performed fluorescent cell-wall and DNA staining during exponential growth and observed that *BL. nexti* cells were predominantly rod-shaped, frequently forming chains with visible septa and polar cell-wall accumulation, consistent with active division and incomplete daughter-cell separation (**Figure 1E**; **Supplementary Figure 1D**). In contrast, *BL. infantis* and *BL. longum* frequently displayed the Y-shaped morphology characteristic of bifidobacteria^37,38^, a phenotype that was nearly absent in *BL. nexti* (<1% of cells). Together, these concordant phylogenomic, genomic similarity, and phenotypic analyses support the designation of *BL. nexti* as a novel subspecies within *B. longum*.

### Subspecies-specific gene content reveals functional divergence among *B. longum* clades

To deepen our understanding of functional differences between the *B. longum* subspecies, we examined subspecies-specific gene content. We screened all 515 isolates’ genomes and 705 MAGs for UniRef90 gene families and quantified their presence across genomes (**Supplementary Figure 2**). Using this pangenome, we applied Principal Component Analysis (PCA) to explore genomic diversity patterns among genomes (**Figure 2A**). We observed four clearly delineated groups, matching our proposed four *B. longum* subspecies. This distinct grouping supports *BL. nexti* has a unique gene repertoire, distinct from other *B. longum* subspecies.

**Figure 2:**
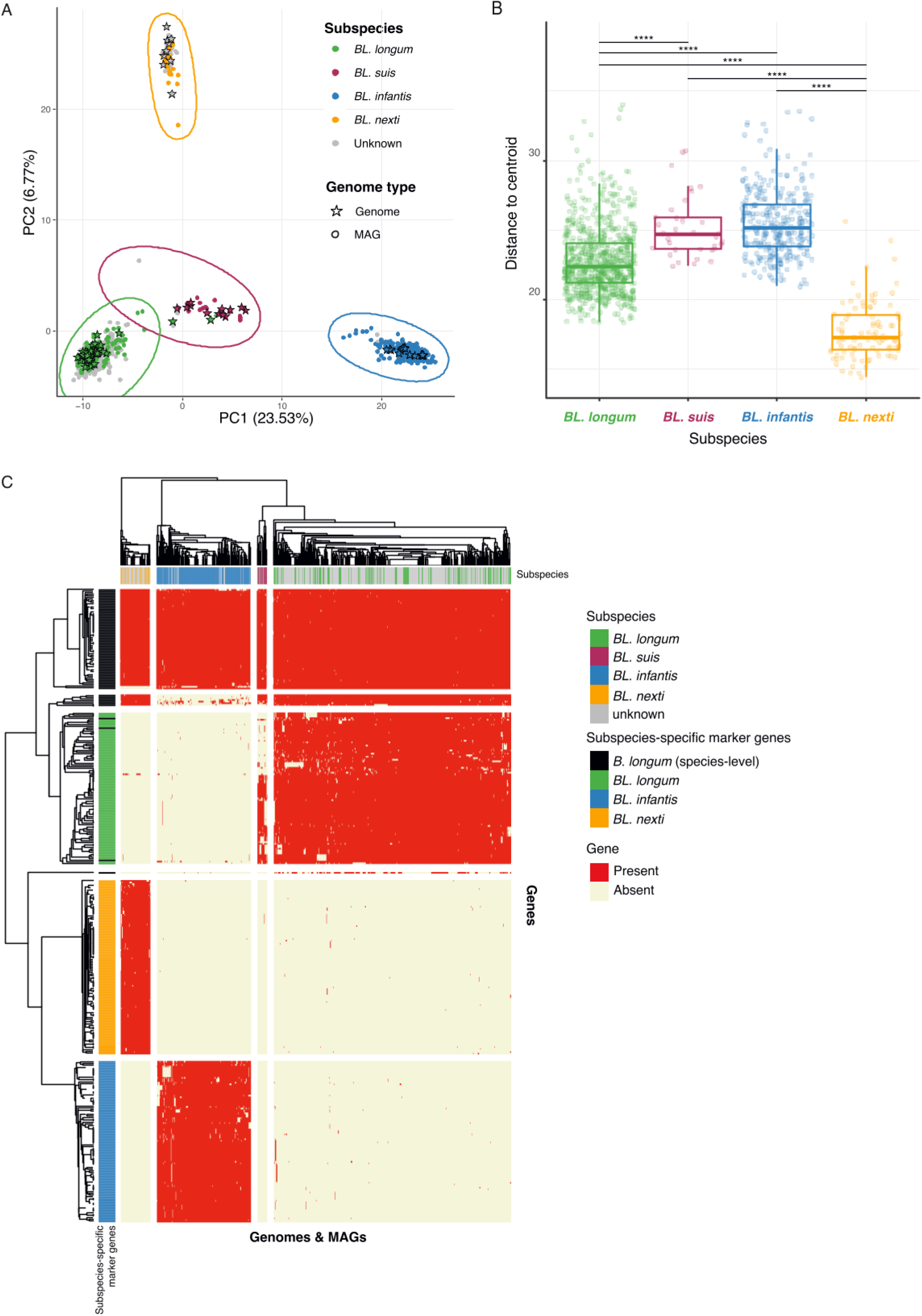
Pangenome analysis identifies subspecies-specific gene content across the four *Bifidobacterium longum* subspecies. **(A)** Principal Component Analysis (PCA) of all *B. longum* genomes (stars) and MAGs (points) based on gene content (presence/absence). Genomes and MAGs are partitioned into four groups (k-means clustering, k=4), with each resulting cluster corresponding to a defined *B. longum* clade. Genomes are colored based on prior public subspecies annotations, while MAGs from the LLNEXT cohort are colored according to their sample annotation as defined in the phylogenetic tree in Figure 1A. **(B)** Boxplot of gene-content variability within each subspecies, based on pangenome gene presence-absence profiles. Variability is calculated as the euclidean distance of each genome/MAG from the centroid of its respective subspecies. To control for unequal sample sizes, analyses were repeated using random subsampling of 34 genomes per subspecies (matching the size of the smallest group, *BL. suis*) over 1,000 iterations, and results were averaged across iterations. ****: *P* <= 0.0001. **(C)** Presence of *B. longum* subspecies-specific and species-level marker genes across all genomes and MAGs. The profiles represent clustering based on Euclidean distances of gene presence/absence. Genes are annotated according to the subspecies for which they are diagnostic, and include the original species-level marker genes from MetaPhlAn 4 (depicted in black), some of which are exclusively found in *BL. longum*.

The internal genomic variability differed significantly across groups (Kruskal-Wallis, P < 2.2×10⁻¹⁶), with *BL. nexti* showing significantly lower variability than all other subspecies (Wilcoxon rank-sum test, P < 2×10⁻¹⁶, **Figure 2B**), suggesting it represents a tighter, more genetically coherent clade. *BL. infantis* showed significantly higher variability than both *BL. nexti* and *BL. longum* (Wilcoxon rank-sum test, P < 2×10⁻¹⁶), likely reflecting its evolutionary specialization for HMO utilization. The diverse metabolic pathways required to process these complex substrates may consequently expand its genomic range. *BL. suis* also showed higher variability than *BL. longum* (Wilcoxon rank-sum test, P = 3.1×10⁻⁷), perhaps since our *BL. suis* definition encompasses *BL. suillum* and *BL. iuvenis*, while this wider genomic diversity can also reflect its transitional niche during weaning^17,18^. Distances to each subspecies’ type strain yielded consistent results (**Supplementary Figure 3A**), with uniform distributions and few outliers across all subspecies, confirming that most isolates relate similarly to their subspecies’ reference type strain.

Next, to examine genome structure conservation and variation across *B. longum* subspecies, we performed whole-genome synteny analysis using the type strain from each clade. We revealed that *BL. nexti* exhibits a high level of genomic similarity to the other *B. longum* subspecies (**Supplementary Figure 3B**). However, we observed a few notable genomic inversions between subspecies, reflecting apparent genome plasticity of the *B. longum* genome^36^. Interestingly, *BL. infantis* was shown to contain genomic regions that are absent from genomes of the other subspecies, consistent with its overall larger genome size.

Defining subspecies-specific genes can help identify key functional traits that distinguish lineages and explain potential ecological or physiological adaptations. Therefore, we searched for UniRef90 gene families that were unique to each subspecies in our pangenome analysis (**Supplementary Figure 2**). We defined a gene as subspecies-specific when it was present in ≥90% of genomes belonging to a given subspecies and in <1% of genomes from each of the other subspecies (**Figure 2C**). Since *BL. longum* and *BL. suis* showed a high amount of gene sharing and clustered close together in the pangenomic space (**Figure 2C, Supplementary Figure 2**), we created subspecies-specific genes only for *BL. longum* without excluding genes shared with *BL. suis*. In total, we identified 59 subspecies-specific genes for *BL. longum*, 77 genes for *BL. infantis*, and 76 genes for *BL. nexti* (**Figure 2C, Supplementary Table 5**).

### *BL. nexti* grows on starch possibly due to a unique α-amylopullulanase gene

To better understand metabolic differences between the *B. longum* subspecies, we focused on genes encoding carbohydrate-active enzymes (CAZymes), which are responsible for degrading, modifying, or synthesizing glycosidic bonds. These enzymes are particularly relevant since each *B. longum* subspecies is known to occupy a distinct ecological niche and specialize in the utilization of different carbohydrate sources. We screened all genomes and MAGs for CAZyme-encoding genes and identified subspecies-specific patterns (**Figure 3A**). In *BL. infantis*, we found several known enzymes associated with the utilization of fucosylated human milk oligosaccharides (HMOs), including GH151 (α-L-fucosidase) and GH95 (exo-α-1,2-L-fucosidase; **Figure 3A**), which are consistent with its specialized metabolic niche. In *BL. longum*, we identified multiple GH43 family enzymes (**Figure 3A**) capable of hydrolyzing arabinan and xylan, polysaccharides derived from plants. These findings reinforce previous evidence that *BL. longum* specializes in plant-derived oligosaccharides^39^. Some of these enzymes were also present in *BL. suis*, suggesting that this subspecies may share partial ability to utilize plant carbohydrates.

**Figure 3:**
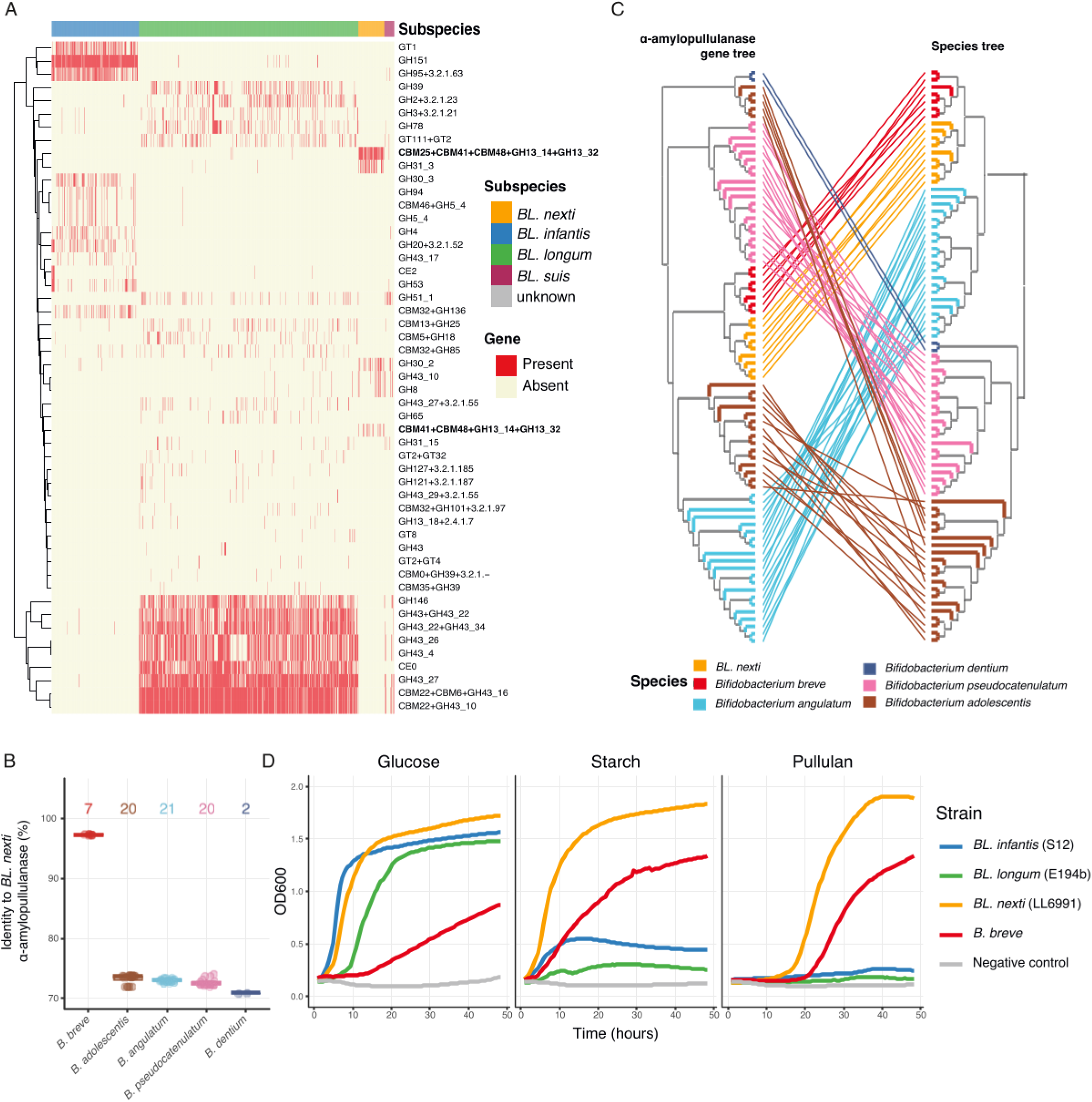
Metabolic profiling and evolutionary history of α-amylopullulanase in *Bifidobacterium longum* subspecies. **(A)** Presence of carbohydrate-active enzymes (CAZymes) across all genomes and MAGs. Genomes are categorized according to their pangenome-based clusters (from Figure 1B), highlighting genes with subspecies-specific distributions. **(B)** Percent identity of the α-amylopullulanase gene across various *Bifidobacterium* species compared to the *BL. nexti* type strain (LL6991) sequence. The number of genomes analyzed per species is indicated at the top. **(C)** Tanglegram comparing the α-amylopullulanase gene phylogeny with the host species tree. Left: nucleotide-based α-amylopullulanase gene phylogeny. Right: corresponding rooted species tree inferred using GTDB-Tk. Lines connect identical genomes between the two trees; tip colors represent GTDB v220 species assignments. **(D)** Growth curves (OD600) of *BL. infantis*, *BL. longum*, *BL. nexti*, and *B. breve* type strains over 48 hours in media supplemented with glucose, starch, or pullulan. Each curve represents the mean of triplicates, with negative controls for each carbohydrate included.

In *BL. nexti*, we detected a unique CAZyme profile, highlighting a distinct carbohydrate utilization strategy (**Figure 3A**). In particular, we identified a single CAZyme gene uniquely enriched in *BL. nexti*, present in 100% of *BL. nexti* genomes, but rare in *BL. suis* (2.94% of genomes), *BL. longum* (0.39% of genomes), and absent in *BL. infantis*. This gene was found to be a *BL. nexti*-specific marker gene as described earlier (**Figure 2C**, bold). This gene encodes two GH13 family domains, GH13_14 (pullulanase; EC 3.2.1.41) and GH13_32 (α-amylase; EC 3.2.1.1), indicating a predicted bifunctional amylopullulanase named α-amylopullulanase, capable of hydrolyzing both (1→6) and (1→4) α-D-glucosidic bonds. Since starch contains both linkage types, this enzyme may enable *BL. nexti* to utilize starch and its components amylose and amylopectin. The gene also included carbohydrate-binding modules CBM41, CBM48, and in most genomes CBM25, which are known to bind α-glucans such as amylose, amylopectin, and pullulan. In addition, it contained a signal peptide (Sec/SPI) and a LPXTG cell wall anchoring motif. Together, these results highlight clear CAZyme specialization across subspecies and support *BL. nexti* as a metabolically distinct subspecies with possible unique starch-utilizing capabilities.

The presence of α-amylopullulanase gene may represent a key evolutionary adaptation for niche colonization within the mammalian gut. To understand the broader phylogenetic distribution of this gene and identify potential horizontal gene transfer events across the genus, we screened 11,141 *Bifidobacterium* genomes cultured from the Dutch Microbiome Project (DMP)^40^. We identified homologs of the α-amylopullulanase in 80 genomes spanning five additional *Bifidobacterium* species (**Figure 3B**, **Supplementary Table 6**). The closest homolog exhibited 97.4% sequence identity to a previously characterized dual-activity α-amylopullulanase from *Bifidobacterium breve*^41^. While we found a highly conserved region within *B. breve*, the gene showed lower similarity to homologs in other human gut-associated *Bifidobacterium* species, including *B. adolescentis*, *B. angulatum*, *B. dentium*, and *B. pseudocatenulatum* (**Figure 3B**, **Supplementary Figure 4A**). As expected, the gene was absent in all *B. longum* genomes that do not belong to the *BL. nexti* subspecies. Comparative synteny analysis of the flanking regions in *BL. nexti* and *B. breve* revealed a homologous 11kb cluster consisting of six genes (**Supplementary Figure 4B**).

The high sequence similarity of the α-amylopullulanase gene between *BL. nexti* and *B. breve*, alongside the presence of a conserved gene cluster prompted us to investigate its evolutionary origin. We therefore compared a GTDB-Tk-based species phylogeny with a phylogeny reconstructed from the α-amylopullulanase gene sequence (**Figure 3C**). In both phylogenetic trees, *B. breve* represented the closest relative of *BL. nexti*. Furthermore, α-amylopullulanase sequences from *BL. nexti* formed a well-supported monophyletic clade, indicating that the gene has evolved cohesively within this subspecies rather than reflecting recent acquisition from *B. breve*. Two evolutionary scenarios are consistent with these observations. First, the gene cluster may have been present in an ancestral *B. longum* lineage and subsequently lost in most descendant lineages while being retained under strong purifying selection in *BL. nexti*. Alternatively, the cluster may have been acquired by an ancestor of *BL. nexti* and subsequently inherited vertically, also under strong purifying selection. To test these hypotheses, we performed a maximum parsimony reconstruction, which inferred that the α-amylopullulanase gene was present in the most recent common ancestor of *B. breve* and the *B. longum* lineage, however the gene was subsequently lost from the ancestral *B. longum* lineage and later reacquired in the ancestor of *BL. nexti* (**Supplementary Figure 4C**). This evolutionary pattern is consistent with horizontal gene transfer (HGT), potentially involving transfer from a *B. breve* lineage, followed by vertical inheritance and conservation within *BL. nexti*.

Previous studies of carbohydrate utilization across *Bifidobacterium* species have shown that growth on starch and pullulan is largely restricted to *B. breve*^42^. To determine whether the α-amylopullulanase gene enables *BL. nexti* to metabolize starch and related polysaccharides as well, we performed growth assays comparing *BL. nexti* with *BL. longum*, *BL. infantis*, and *B. breve*. As expected, *BL. longum* and *BL. infantis* showed minimal growth on starch and pullulan, whereas *B. breve* grew well on both substrates, validating the experimental conditions (**Figure 3D**). In contrast, *BL. nexti* exhibited robust growth on starch and pullulan, reaching an OD_600_ exceeding 1.5 (**Figure 3D**). This phenotype was confirmed in two additional *BL. nexti* isolates, both of which grew on starch, although one failed to utilize pullulan, suggesting some strain-level variability in substrate utilization (**Supplementary Figure 5A**). Growth was also observed on glycogen in an additional assay (**Supplementary Figure 5B**), further supporting the ability of *BL. nexti* to utilize branched α-glucans. Consistent with these findings, RT-qPCR revealed a 1.6-fold increase in α-amylopullulanase gene expression during growth on starch compared with glucose (**Supplementary Figure 5C**). Together, these results suggest that *BL. nexti* occupies a metabolic niche distinct from other *B. longum* subspecies by enabling the utilization of starch-derived polysaccharides that become available in the infant gut during dietary transition.

### *B. longum* subspecies dynamics in mother-infant dyads from pregnancy to post-partum

Next, we utilized subspecies-specific marker genes to quantify the metagenomic abundance of three *B. longum* subspecies (*BL. longum*, *BL. infantis*, and *BL. nexti*; no subspecies-specific marker genes could be determined for *BL. suis*, **Methods**) within the LLNEXT birth cohort^31^. Previous investigations targeted the maternal stool metagenomes as a potential reservoir and source of infant strains^7,43,44^. As expected, *BL. longum* was highly prevalent in maternal samples across all time points (prevalence 94–96.4%), though at low relative abundance (**Figure 4A, Supplementary Figure 6A**). Detection of *BL. infantis* in mothers was rare and at very low relative abundance (<0.1%, **Figure 4A**). Only one of 414 mothers (0.24%) carried *BL. infantis* at gestational week 12, increasing slightly to 4/406 and 5/568 mothers (1–1.87%) at week 28 and at delivery, respectively, and decreasing to 3/499 mothers (0.6%) at three months postpartum (**Figure 4A**). Important to note, some of these gains and losses were impacted by slight differences in the mothers sampled over time. *BL. nexti* was detected more frequently in mothers, appearing in 2.4% of mothers at week 12 and increasing to 4.1% around delivery, though also consistently below 1% relative abundance. Notably, *BL. nexti* was not detected 3 months postpartum, suggesting transient carriage around delivery (**Figure 4A**). *BL. nexti* strains carried by infants were more similar to their mothers than to unrelated individuals (marker gene major allele identity, Wilcoxon signed-rank test, *P*=0.0046, **Methods**), consistent with vertical transmission but not definitive. Overall, these results indicate that while *BL. longum* is commonly present in mothers, *BL. infantis* is exceedingly rare, supporting the conclusion that infant colonization with *BL. infantis* likely originate from sources other than maternal gut. However, we acknowledge the possibility that *BL. infantis* may be present in mothers below metagenomic detection limit at this sequencing depth. In contrast, *BL. nexti* increases during pregnancy and is more frequently detected around the time of delivery, suggesting it may be a potential source transmitted from mothers to infants during or shortly after birth.

**Figure 4:**
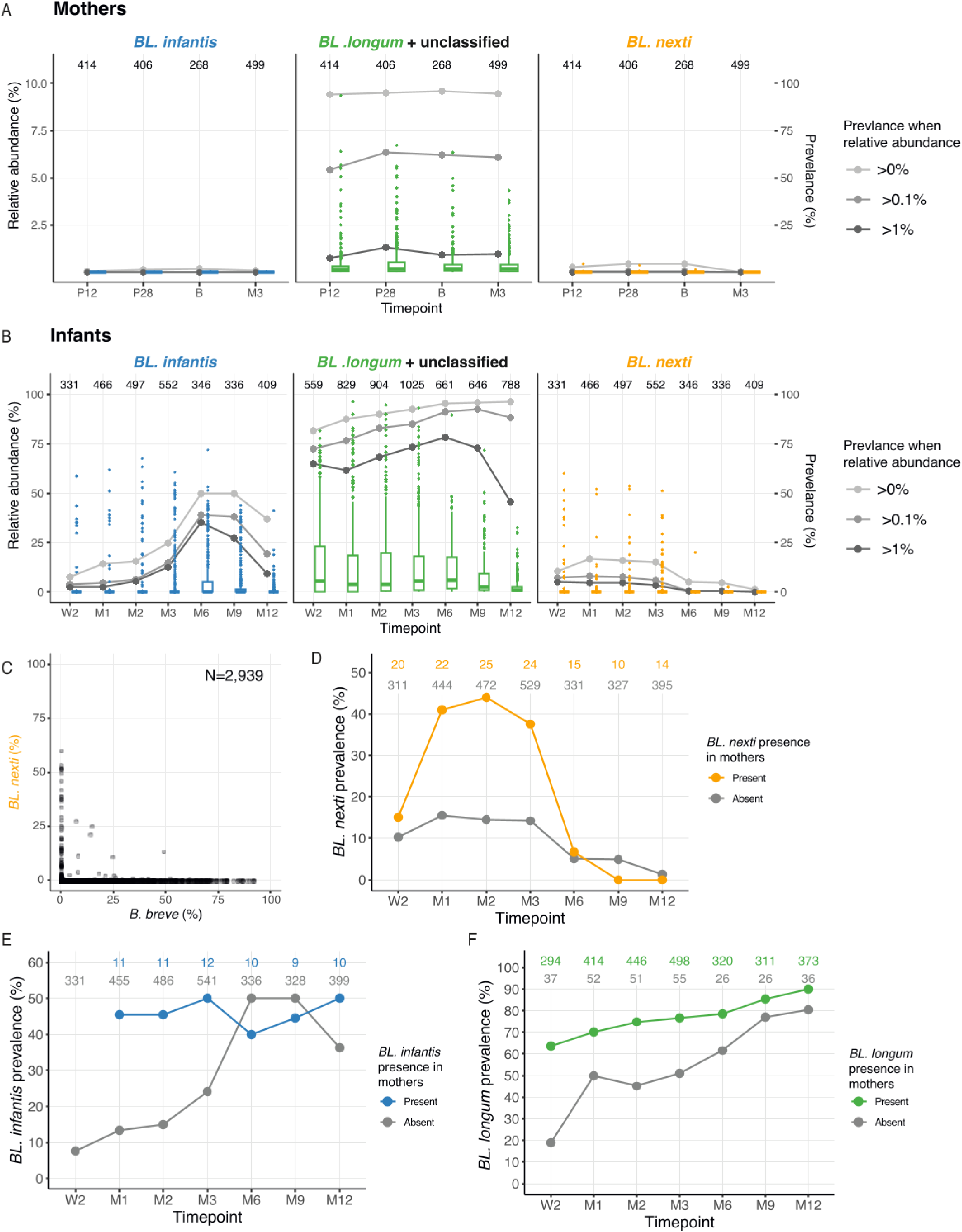
*B. longum* subspecies dynamics within the LLNEXT cohort. **(A-B)** Boxplots showing the longitudinal relative abundance of *BL. infantis*, *BL. longum* together with unclassified *B. longum* and *BL. nexti* in **(A)** mothers and **(B)** infants. Overlaid lines represent subspecies prevalence calculated across various relative abundance thresholds. The absolute number of individuals (N) sampled at each time point is indicated at the top of each panel. **(C)** Correlation between *B. breve* and *BL. nexti* relative abundance across all samples. Number of samples analyzed shown. **(D-F)** Longitudinal prevalence of *B. longum* subspecies in infants, stratified by maternal colonization status (mothers who carried the subspecies at any time point versus mothers who did not) for **(D)** *BL. nexti*, **(E)** *BL. infantis*, and **(F)** *BL. longum*. Presence of a subspecies in mothers was associated with higher prevalence in infants.

To further investigate whether *B. longum* subspecies in the infant gut originate from other, non-gut maternal sources, we analyzed 82 maternal vaginal samples using metagenomic data. We observed that *BL. nexti* was present in 6% (5/82) of the maternal vaginal samples, with relative abundances ranging from 0.02% to 4.2%. No other subspecies of *B. longum* were found in vaginal samples. To further validate this result, we used PCR with *BL. nexti*-specific primers in two specific samples (with higher *BL. nexti* relative abundance, 0.2% and 4.2%), which resulted with positive *BL. nexti* presence in both samples. Interestingly, *BL. nexti* was not detected in the gut metagenomes of these specific mothers nor was it detected with PCR amplification. Maternal vaginal carriage of *BL. nexti* was associated with an increased trend in the odds of infant colonization at any time point (3 infants out of 5 mothers vs. 158 infants out of 680 mothers; generalized linear model, log-odds_ever_present_=1.6, *P* = 0.08).

Next, we examined the dynamics of *B. longum* subspecies colonization in infants from birth through the first year of life (**Figure 4B**). *BL. longum* was the most prevalent subspecies throughout infancy, increasing from 81% prevalence at two weeks of age to 96% at one year (**Figure 4B, Supplementary Figure 6B**). However, despite its high prevalence, median relative abundance of *BL. longum* declined by one year, indicating that many infants carried it only at low levels. In contrast, *BL. infantis* was less prevalent during early infancy than its established reputation as a dominant gut symbiont might suggest; only 15% of infants exhibited detectable levels within the first two months of life (**Figure 4B**), consistent with previous observations in other westernized cohorts^30^. Its prevalence increased over time and peaked at six months, however even then only 50% of infants carried *BL. infantis*, and just 35% exceeded 1% relative abundance (**Figure 4B**). These patterns reinforce reports of the progressive disappearance and competitive decline of *BL. infantis* in westernized populations^45,46^. *BL. nexti* showed distinct colonization dynamics. It was primarily detected in the early months of life, beginning at 10.6% prevalence at week 2 and rising to 15.2-16.7% between one and three months (**Figure 4B**). Relative abundances were generally low; while a small number of infants exhibited high levels (>10% relative abundance), the majority carried *BL. nexti* only transiently and at low abundance (**Figure 4B**). This pattern suggests that *BL. nexti* may represent an early-life colonizer with limited long-term persistence in the developing gut, potentially shaped by competition with other *Bifidobacterium* species or *B. longum* subspecies.

Given that *BL. nexti* and *B. breve* share the ability to utilize starch carbohydrates (**Figure 3D**), we investigated their co-occurrence patterns to determine if they compete for these substrates within the infant gut environment. We observed a strong and clear negative association between the presence of *BL. nexti* and *B. breve* (effect_abundance_=-0.0133, *P*=1.81x10^-3^, **Figure 4C, Supplementary Table 7**). This finding is consistent with a hypothesis of species competition, possibly driven by shared metabolic niches and competition for specific complex carbohydrates in the infant gut environment. Furthermore, *BL*. *nexti* was rarely found to co-occur with *BL*. *infantis* or *BL. longum* (**Supplementary Figure 6D,E**), suggesting both interspecies and intraspecies competition.

Finally, we assessed whether infants harboring specific *B. longum* subspecies were more likely to have mothers carrying the same subspecies in their gut. We observed a significant association between the presence of *BL. nexti* in the gut of mothers and their infants (**Figure 4D**). Maternal carriage of *BL. nexti* was a significant predictor of both increased abundance in early life (up to three months; effect_abundance_=1.37, *P*=0.018) and the likelihood of infant acquisition at any point during the study period (log-odds_ever_present_=1.35, *P*=4.3x10^-4^). For *BL. infantis*, although maternal carriage was very rare, mothers harboring the subspecies tended to have infants with significantly higher *BL. infantis* abundance (effect_abundance_=2.1, *P*=0.03; **Figure 4E**). Similarly, the presence of *BL. longum* in mothers was strongly associated with higher abundance in their offspring (effect_abundance_=3.05, *P*=9.29x10^-7^; **Figure 4F**). Collectively, these findings support the hypothesis that vertical transmission from the maternal gut or vaginal microbiome is a plausible source for the seeding of *BL. nexti* and other *B. longum* subspecies in the infant gut.

### Association of *B. longum* subspecies with early-life phenotypes

To better understand the possible drivers of *B. longum* subspecies presence and abundance, and the physiological consequences of this taxon in early life, we conducted multivariable associations with each *B. longum* subspecies and information on delivery, feeding and parity available in LLNEXT cohort^31^. The abundance of *BL. infantis* in infants was not associated with delivery mode, delivery place or parity. Interestingly formula-fed infants showed a higher presence of *BL. infantis* (log-odds_ever_present_=0.71, *P*=4.9x10^-4^, **Figure 5A, Supplementary Table 7**). Early life abundance of *BL. longum* in infants was higher in infants delivered vaginally (effect_abundance_=2.04, *P*=8.23x10^-5^, **Figure 5B**), and within the group of vaginally delivered infants, higher in home vs hospital delivery (effect_abundance_=1.47, *P*=1.13x10^-3^, **Supplementary Figure 7A**). In addition, *BL. longum* was higher in formula-fed infants (effect_abundance_=0.774, *P*=2.21x10^-2^, **Figure 5C**) and in non-firstborns (effect_abundance_=1.68, *P*=4.58x10^-5^, **Supplementary Figure 7B**). *BL. nexti* abundance was negatively associated with infant age (effect_abundance_=-0.005, *P*=2.93x10^-32^, **Figure 4B**). We evaluated the impact of infants’ delivery mode and found that *BL. nexti* had a significantly higher abundance and was more prevalent in infants delivered via C-section (effect_abundance_=1.17, *P*=3.19x10^-4^; log-odds_ever_present_=-0.95, *P*=5x10^-5^, **Figure 5D**). In vaginally-delivery infants, *BL. nexti* abundance did not show a strong relationship with either home or hospital delivery (**Supplementary Table 7**). In contrast, the presence of older siblings was negatively associated with *BL. nexti* abundance and prevalence (effect_abundance_=-0.65, *P*=2.76x10^-4^; log-odds_ever_present_=-0.45, *P*=0.04, **Supplementary Figure 7C**).

**Figure 5:**
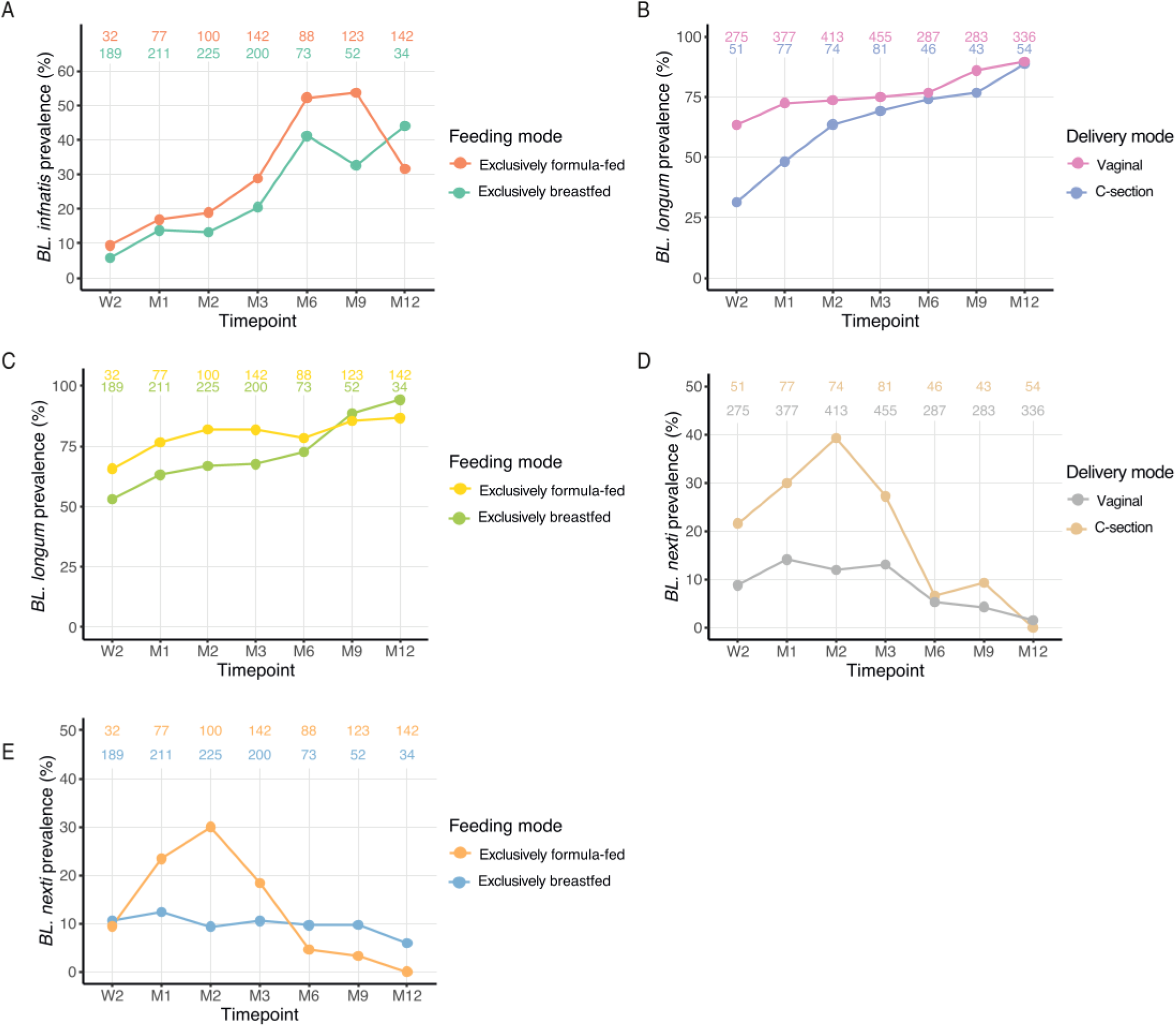
Association of early-life factors with the presence of *B. longum* subspecies in the infant gut microbiome. **(A-E)** Longitudinal prevalence of *B. longum* subspecies in infants stratified by early-life factors. Plots show the prevalence over time for **(A)** *BL. infantis*, **(B-C)** *BL. longum*, and **(D-E)** *BL. nexti*. Infants are stratified by **(A,C,E)** feeding mode (exclusively breastfed versus exclusively formula-fed) and **(B, D)** delivery mode (vaginal delivery versus C-section). Absolute numbers for each group (N) are indicated at the top of each plot.

Given the diverse carbohydrate utilization profiles of *BL. nexti* compared to othe*r B. longum* subspecies, we investigated the potential influence of feeding mode at each timepoint on their prevalence in the infant gut. At early timepoints (week 2) no differences were observed between infants who were exclusively formula-fed and infants who were exclusively breast-fed (**Figure 5E**). However, *BL. nexti* abundance in formula-fed infants increased over time, reaching a maximum prevalence at month 2, while breast-fed infants did not increase overtime (likelihood ratio test (LRT) feeding mode-time interaction, *P*=1.35x10^-5^; log-odds_ever_present_ =0.73, *P*=2.5x10^-3^; **Figure 5E**). This indicates a possible selective advantage of formula feeding promoting the growth of *BL. nexti* strains, possibly due to the starch utilization ability of *BL. nexti*. Beyond three months of age, no differences were observed in the prevalence of *BL. nexti* between formula-fed and breast-fed infants.

### *BL. nexti* is present globally at low relative abundance

To assess the global distribution and dynamics of *BL*. *nexti*, we examined its prevalence across multiple early-life international cohorts and stratified samples into four early-life age bins (0-1, 1-3, 3-6, and 6-12 months). This analysis included 6,162 samples from 18 locations across the world. *BL. nexti* was detectable in infants from many geographic regions but generally at very low relative abundance (**Figure 6A**). Its prevalence ranged from nearly 100% in early infancy, as seen in Oregon, USA to lower prevalences found in Georgia, USA, Luxembourg and the Netherlands. Notably, its temporal pattern varied substantially between cohorts. In several locations, prevalence was highest in early infancy, as observed in the LLNEXT cohort, yet in many other regions *BL. nexti* showed the opposite trend, with higher detection rates at later time points (**Figure 6A**). Only a minority of infants had *BL. nexti* which exceeded a relative abundance of 0.1% (**Figure 6A**, filled boxes), indicating that while the subspecies is globally widespread, it typically remains a low-abundance member of the infant gut microbiome. These patterns collectively support *BL. nexti* as an early-life associated bifidobacterial lineage with modest but detectable prevalence across populations.

**Figure 6:**
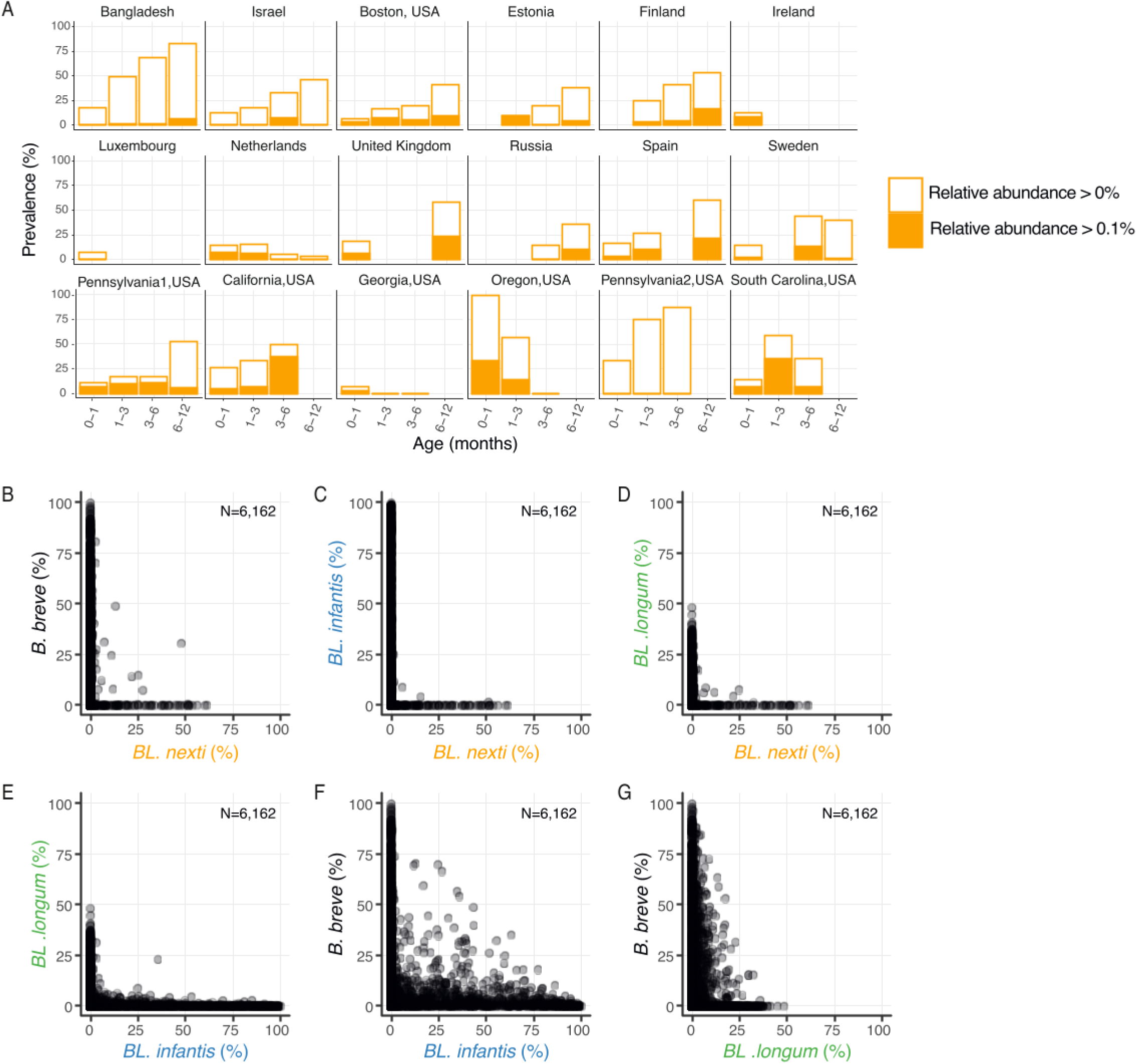
Global prevalence and co-occurrence patterns of *BL. nexti.* **(A)** Global prevalence of *BL. nexti* across international cohorts, categorized by infant age bins (0-1, 1-3, 3-6, and 6-12 months). Prevalence is reported using two detection thresholds: relative abundance >0% and >0.1%. **(B-G)** Correlation analysis between the relative abundance of *BL. nexti* and other *Bifidobacterium* taxa, including *B. breve* and additional *B. longum* subspecies, across all infant samples.

Given that *BL. nexti* and *B. breve* share the metabolic capacity to utilize starch-like substrates, and considering that subspecies of the same species often occupy overlapping ecological niches, we extended the co-occurrence analysis initially performed on the LLNEXT cohort to all analyzed cohorts. Our global analysis revealed a prominent pattern of mutual exclusion between *BL. nexti* and *B. breve*, so that *BL. nexti* rarely achieved high relative abundance in samples where *B. breve* was also detected (**Figure 6B**). When *B. breve* was present, *BL. nexti* abundance was almost always below 1%, strongly suggesting a mechanism of competitive exclusion or significant niche overlap between these two lineages. A similar pattern of mutual exclusivity was also observed between *BL. nexti* and *Bifidobacterium adolescentis* (**Supplementary Figure 8**), although this species is less relevant in the infant gut context due to its low prevalence in infants and its primary association with the adult microbiome.

This mutually exclusive pattern also extended to subspecies within the *B. longum* species. We found that the presence of either *BL. infantis* or *BL. longum* was also associated with the low abundance of *BL. nexti* (**Figure 6C,D**). While *BL. nexti* was never abundant in samples containing *BL. infantis*, its prevalence was predominantly at low levels (<1%) in samples with *BL. longum*. Interestingly, *BL. longum* and *BL. infantis* themselves exhibited largely mutually exclusive patterns, with most samples being dominated by only one of the two subspecies (**Figure 6E**). However, in contrast to previous reports^47^, we observed the co-occurrence of *B. breve* and *BL. infantis* (**Figure 6F**), particularly across larger cohorts (in Bangladesh, the Netherlands, and Boston, USA). In addition, we found co-occurrence of *BL. longum* and *B. breve* (**Figure 6G**), both known to be highly prevalent in infants. Collectively, these findings demonstrate that *BL. nexti* generally fails to establish high-abundance colonization when competing subspecies or *B. breve* are present. This reinforces the concept that the infant gut community typically supports only a single dominant *B. longum* subspecies, with *BL. nexti* occupying a limited and highly competitive ecological niche.

## Discussion

In this study, we performed a comprehensive genomic, phenotypic, and phylogenomic characterization of the *Bifidobacterium longum* species complex, robustly establishing *B. longum* subsp. *nexti* as a distinct, novel subspecies. Supported by multiple complementary genomic approaches, including average nucleotide identity (ANI), digital DNA-DNA hybridization (dDDH), and pangenome analysis, this classification aligns with modern polyphasic prokaryotic taxonomy that integrates traditional methods with advanced genome-based analyses. Beyond taxonomic resolution, we developed subspecies-specific marker genes for metagenomic quantification and characterized the global dynamics of *BL*. *nexti*, revealing its widespread presence at low relative abundance. Furthermore, we identified a unique ecological niche for this subspecies centered on starch degradation, and noted its mutual exclusivity with *B. breve* and other *B. longum* subspecies within the infant gut, suggesting a distinct competitive landscape.

Defining rigid taxonomic boundaries within the *B. longum* species remains a significant challenge due to the limited genetic diversity and high degree of homology between lineages, which likely represent groups in a state of relatively recent evolutionary divergence. The high amount of gene sharing, particularly the close clustering and substantial gene content overlap between *BL*. *suis* and *BL*. *longum*, prevented our MetaPhlAn-based approach from resolving *BL*. *suis* as a distinct clade. Critically, a substantial proportion of metagenomic sequences belonging to the *B. longum* species remained "unclassified" within our metagenomic analysis. While some of this is due to subspecies overlap, the prevalence of these unclassified sequences suggests that much of the diversity within the complex remains hidden and that the current taxonomic framework is not yet exhaustive. The identification of *BL. nexti* suggests that as sequencing depth and cohort sizes increase, additional niche-specific subspecies will likely be uncovered from the remaining reservoir of unclassified *B. longum* diversity.

This study has several limitations that also highlight important opportunities for future research. Although comparative genomics, growth assays, and increased α-amylopullulanase expression during growth on starch strongly support a role for this locus in starch utilization, definitive confirmation will require targeted gene knockout and complementation experiments. While our analysis included more than 6,000 metagenomes from geographically diverse cohorts, expanding to additional non-Westernized populations will further refine our understanding of the global distribution and ecological significance of *BL. nexti*. Because *BL. nexti* was generally present at low relative abundance, deeper metagenomic sequencing and targeted detection approaches may provide a more complete picture of its colonization dynamics. Finally, although the ecological and epidemiological associations identified here were robust across this large dataset, they remain correlative and will benefit from validation in independent cohorts and prospective experimental studies to establish the mechanisms underlying vertical transmission, pregnancy-associated enrichment, and diet-related selection.

The early-life presence of *BL. nexti* presents an intriguing ecological puzzle given its apparent capacity for starch degradation. We initially hypothesized that this subspecies would emerge later in infancy, coinciding with the introduction of solid foods. However, in the Dutch LLNEXT cohort, *BL. nexti* prevalence peaked between one and three months of age, a period during which dietary starch is largely absent, particularly in exclusively breastmilk-fed infants. One possible explanation for its early appearance is that infant colonization may not depend solely on dietary exposure, but may instead reflect transmission from maternal microbial reservoirs. A recent study identified *BL. nexti* in the oral microbiome^48^, where its starch-degrading capacity could provide a selective advantage in the presence of dietary starch. Maternal oral colonization may therefore serve as a source of early-life acquisition, a possibility supported by the higher prevalence of *BL. nexti* among formula-fed infants, which exhibit greater mother-infant oral microbiome sharing compared with breastfed infants. Another possibility is that the α-amylopullulanase gene provides a selective advantage by enabling the utilization of substrates other than dietary starch. One plausible candidate is glycogen, which shares the α-1,4- and α-1,6-glycosidic linkages characteristic of starch and pullulan. Here, we showed that *BL. nexti* is also capable of growing on glycogen. Glycogen is present in breast milk and can additionally be produced endogenously by the infant from maltodextrin and lactose-derived carbon sources, providing a potential early-life niche for this metabolic capability. Another option can be explained by the observed mutual exclusivity between *BL. nexti*, *B. breve*, and other *B. longum* subspecies. This suggests strong niche competition within the early-life gut ecosystem. Given that *B. breve* often dominates infant colonization, the persistence of *BL. nexti* likely depends on specialized functional traits beyond carbohydrate utilization alone. These may include unique adhesion factors, competitive exclusion mechanisms, or distinct interactions with the host immune system, all of which remain to be elucidated.

Beyond the infant gut, we observed an increased prevalence of *BL. nexti* in mothers during late pregnancy. This pattern is consistent with previous reports showing that progesterone levels modulate the composition of *Bifidobacterium* species during pregnancy^49^. In the vaginal environment, progesterone promotes glycogen deposition by epithelial cells, thereby creating a carbohydrate-rich niche that supports bifidobacterial growth^50^. This raises the possibility that *BL. nexti* may also be adapted to persist in the vaginal niche, potentially enhancing vertical transmission and increasing the likelihood of neonatal colonization by bifidobacteria more broadly. The detection of *BL*. *nexti* in 6% of maternal vaginal samples, which we initially identified using our adapted MetaPhlAn-based method and subsequently validated via PCR, suggests vertical transmission as a potential colonization source. This successful cross-method validation not only confirms the presence of the subspecies in the vaginal niche but also reinforces the sensitivity and accuracy of our computational approach.

On a global scale, we found that *BL. nexti* is widespread but typically occurs at very low relative abundance, often below 0.1%. Such low abundance likely reduces detection sensitivity in metagenomic datasets, particularly in smaller cohorts or datasets with limited sequencing depth, thereby limiting the statistical power to detect ecological and clinical associations. We anticipate that deeper metagenomic sequencing and the inclusion of more diverse populations will reveal additional *BL. nexti* colonization events. Consistent with this, ultra-deep sequencing studies of the human oral microbiome have reported a substantial prevalence of *BL. nexti*^48^, suggesting that increased sequencing depth can improve its detection. Its persistently low abundance in the infant gut may also reflect strong ecological competition with *B. breve* and other *B. longum* subspecies.

Despite its rarity, the consistent detection of *BL. nexti* across multiple geographic regions and body sites, including the oral cavity, gut, vaginal microbiome, and breast milk, highlights an unusual multi-site carriage pattern. This broad ecological distribution appears to be uncommon among early-life-associated bifidobacteria and suggests that *BL. nexti* occupies a distinct ecological niche spanning maternal and infant-associated environments. Together with its specialized metabolic repertoire and enrichment in non-gut environments observed in the *B. longum* subspecies phylogeny, these findings further support its designation as a novel *B. longum* subspecies. Despite its low abundance, the consistent detection of *BL. nexti* across multiple geographic regions and body sites^20^, including the oral cavity^48^, gut, vaginal microbiome, and breast milk, highlights an unusual multi-site carriage pattern. Together with its specialized metabolic repertoire these findings further support its designation as a novel*B. longum* subspecies.

## Data availability

The type strain of *Bifidobacterium longum* subsp. *nexti*, LL6991, was isolated from an infant enrolled in the LLNEXT cohort. The GenBank accession numbers for the 16S rRNA gene sequence and the draft genome assembly are PV770940 and JBPCCJ000000000, respectively.

The *B. longum* MAGs and genomes analyzed in this study have been deposited in Zenodo https://doi.org/10.5281/zenodo.21280061.

Quality-trimmed, human-decontaminated shotgun metagenomic sequencing data from the LLNEXT cohort are publicly available at the NCBI SRA under accession number PRJNA1468137. Metadata (including questionnaire and clinical information) and quality-trimmed, human-decontaminated sequencing reads are available under accession numbers EGAD50000001187 (fecal samples), EGAD50000002469 (vaginal samples), and EGAD50000002677 (metadata). Access can be requested through the application form available at https://forms.gle/A4Jem2rMnjcygWRD6. Access to Lifelines NEXT data is available to all qualified researchers and is governed by the Lifelines NEXT Data Access Agreement (https://groningenmicrobiome.org/?page_id=2598). This controlled-access procedure ensures that metadata are used exclusively for scientific research purposes and in accordance with the informed consent provided by Lifelines NEXT participants. To facilitate data access and reuse, we have prepared a detailed instruction guide describing the EGA access procedure, available at: https://github.com/GRONINGEN-MICROBIOME-CENTRE/LLNEXT_pilot/blob/main/Data_Ac cess_EGA.md. No fees are associated with access to the metadata or sequencing data.

## Methods

### Data Collection and assembly of MAGs

For this study, we curated a dataset of 1,702 genomes and metagenome-assembled genomes (MAGs) belonging to the *B. longum* species complex. To minimize analytical noise and ensure high-quality genomic analysis, we selected 1,199 genomes with >95% completeness and <5% contamination, as determined by CheckM2 (v1.0.2)^51^. Altogether these included 501 isolate genomes and 698 *B. longum* MAGs. A total of 114 genomes were retrieved from NCBI by searching for entries matching *Bifidobacterium longum*. Similarly, 169 genomes were obtained from the JGI IMG/M (Integrated Microbial Genomes & Microbiomes) database. Subspecies annotations were extracted from strain names when available. In addition, four *B. longum* genomes from Chilean isolates were included from Díaz et al.^52^. Two isolates were isolated as part of the Dutch Microbiome Project (DMP), as described below^40^. 49 genomes were sourced from the Baby Biome Study (BBS) in the United Kingdom^53^, with details on isolation, sequencing and assembly as previously described^53,54^. In addition, 323 MAGs were assembled from this cohort as previously published^53,54^. An additional 99 genomes were obtained from the MicrobeMom study in Ireland, with full methods described therein^55^. 18 MAGs were gathered from the Amsterdam Infant Microbiome Study (AIMS) in the Netherlands^14^. 14 genomes were taken from a study in Bangladesh^19^. 49 genomes were taken from isolates in Spain as part of the MAMI cohort^56^. One isolate strain (LL6991) was obtained from the LLNEXT cohort, as described below. In addition, 346 MAGs were created from this cohort as described previously^57^. For all genomes subspecies annotations were derived from available strain nomenclature. MAGs from samples in the LLNEXT cohort that clustered within the *BL*. *nexti* clade were annotated as *BL*. *nexti*. Otherwise genomes and MAGs were classified as "unknown".

11 MAGs from the Breastmilk Baby (BMB) Israeli cohort (BioProject: PRJNA994433), were created using the following pipeline: Host reads were removed using an in house pipeline by aligning reads to the human genome by Bowtie2^55^ (2.4.5-1). Samples were filtered and trimmed for Nextera adapters using fastq-mcf, ea-utils^56^ (1.05). We performed metagenomic assembly using MEGAHIT (1.2.9)^58,59^ to reconstruct contigs from the raw reads. To facilitate binning, we mapped the original reads back to these contigs using minimap2 (2-2.24)^60^, generating coverage profiles for each assembly. We then performed automated binning with MetaBAT2 (2:2.17)^61^, integrating the assembly files and mapping results to group contigs into discrete genomic bins. To ensure the reliability of the resulting MAGs, we assessed their quality,including completeness and contamination, using CheckM2 (1.0.2)^51^. Finally, we assigned taxonomic classifications to the high-quality MAGs using the Genome Taxonomy Database Toolkit (GTDB-Tk, 2.3.2)^62–64^, and identified members of the *B. longum* species complex.

### Strain isolation, DNA extraction, sequencing and assembly

Two isolates were isolated as part of the Dutch Microbiome Project (DMP)^40^, as follows: we collected stool samples from 550 population-based participants from the Lifelines Dutch Microbiome project. We used two distinct sampling strategies to culture bacterial isolates. In the first round of culturing, we chose selective culturing conditions for specific taxonomic groups, in this caseDe Man–Rogosa–Sharpe (MRS, Millipore, 69964) medium under anaerobic conditions for genus *Bifidobacterium* (phylum Actinobacterium). We assigned species for each isolate via multiplexed 16S sequencing for each isolate and performed whole-genome sequencing (WGS) on a subset (on average two isolates per species per donor). In the second round of culturing, we selected dozens of isolates per donor based on diverse morphological characteristics, cultured all in a rich medium, Brain Heart Infusion (Bacto 237300), and performed WGS. DNA extraction was performed using the PureLink Pro Genomic DNA Purification (K182104A) Kit.

The type strain, LL6991^T^, was isolated in April 2025 in Groningen, Netherlands (53.2210° N, 6.5770° E) from a fecal sample taken from a two-weeks old infant as part of the Lifelines NEXT cohort, a birth cohort designed to study the effects of intrinsic and environmental determinants on health in mothers and children^57^. A detailed description of the design, inclusion and exclusion criteria has been described previously^31^. For isolation, a stool sample that was known to contain the type strain was spread-plated onto Yeast Casitone Fatty acid agar^65^ supplemented with 20 mM Lactose and 0.05% (w/v) cystine (YCFA-Lactose) were then incubated in an anaerobic chamber (90% nitrogen, 5% hydrogen, 5 % carbon dioxide) for 72 hours. Each quadrant of the spread plate was subcultured onto fresh YCFA-Lactose agar plates after which individual colonies were picked up randomly and subcultured on fresh YCFA-Lactose agar for colony purification. These purified colonies were identified up to species level using direct transfer method matrix-assisted laser desorption/ionization coupled to a time-of-flight mass spectrometer (MALDI-TOF smartfleX, Bruker Daltonics, Germany) where one of the isolates obtained was identified as *Bifidobacterium longum.* Further, the DNA of a *B. longum* isolate was extracted using QiAmp DNA Mini kit (Qiagen, Hilden, Germany) and PCR reactions were run using Applied Biosystems™ Amplitaq gold 360 Master mix (Thermo Fisher, USA) and primers that bind to 4 gene loci found to be unique to *B. longum* subsp. *nexti* to verify its subspecies identity (dasR_F: CGAACGAGATTACGGTCTGC, dasR_R: CATGATGCATGGTGAATCGCA, group_10176_F:GTGGTCTGGGCTGTCGAATC, group_10176_R: CGAGAATATCGGTCAGGGGC, group_865_F: GAACCATTACCCCAGCTCCC, group_865_R: CAGGTCATAGGCGAGGCATT, group_12984_F: AATTTATCGCCGCTGGACTT, group_12984_R: GACTTCATCGCTGCTACCACA).

For the type strain, LL6991, and for an additional *BL. nexti* strain, MB0413, DNA was extracted using MagAttract HMW DNA Kit (#67563, Qiagen). Short read sequencing libraries were prepared using Nextera XT DNA Library Preparation kit (FC-131-1096, Illumina) according to the manufacturer’s recommended protocol with half of the volume and the DNA. Samples were sequenced using Illumina single-end 150 bp sequencing on a NextSeq 500 device. Long reads were retrieved using native barcoding sequencing kit (SQK-NBD114-24, ONT) on a MinION device. Basecalling and demultiplexing of long reads was done using Dorado (Oxford Nanopore Technologies). Hybrid assembly was performed using SPAdes (v4.0.0)^66,67^.

For the two strains derived from the UK Baby Biome Study infants, 37822_2_275 and 37822_2_213, hybrid genome assemblies were generated. DNA was extracted using the MasterPure Complete DNA and RNA Purification Kit (Lucigen), then sequenced on the Illumina NovaSeq 6000 (384-plex, 2 x 151bp, Illumina) and on the Oxford Nanopore GridION (24-plex) or PromethION (96-plex) using R9.4.1 and R10.4.1 flow cells with basecalling in ‘super-accurate (SUP) model using Dorado. Hybrid genome assemblies were generated using the Dragonflye with Flye for long-read assembly (‘--nanohq’ with R9.4.1 data and ‘--nano_hq --read-error 0.03’ with R10.4.1 data), Medaka (https://github.com/nanoporetech/medaka) for long-read polishing and Polypolish for short-read polishing.

### Lifelines NEXT (LLNEXT) cohort analysis

Longitudinal gut metagenomes from 714 mother–infant pairs enrolled in the Lifelines NEXT (LLNEXT) cohort were included in this study^57^. In total, 4,526 fecal metagenomes were analyzed, comprising 1,587 maternal samples collected at 12 and 28 weeks of pregnancy, at delivery, and 3 months postpartum, and 2,939 infant samples collected at 2 weeks and at 1, 2, 3, 6, 9, and 12 months of age. Maternal samples yielded a median of 13.7 million paired-end raw reads and 12.8 million paired-end quality-filtered reads per sample, while infant samples yielded a median of 14.7 million paired-end raw reads and 13.8 million paired-end quality-filtered reads per sample.

A marker-gene-based phylogeny of *B. longum* was reconstructed using StrainPhlAn v4.0. Marker genes corresponding to the *B. longum* species-level genome bin (SGB17248) were extracted from maternal and infant fecal samples in the LLNEXT cohort, aligned, and used to infer a phylogenetic tree. Reference genomes (n = 20) representing the four *B. longum* clades were integrated into the phylogeny using the -r flag. Relative abundances of *B. longum* subspecies s (*BL. infantis*, *BL. longum*, and unclassified *B. longum*) were estimated using MetaPhlAn v4.0 with our previously described custom database^30^, which includes subspecies-level marker-genes. Subspecies-level abundances (*BL. infantis*, *BL. longum*, and unclassified *B. longum*) were calculated by normalizing the summed abundance of each group to the total *B. longum* abundance within each sample.

### Genomic, phylogeny and pangenomic analysis

Average nucleotide identity between genomes was estimated using FastANI (1.33)^68^, and mean values were aggregated within each *B. longum* clade.. The selection of the number four for clustering was done using the Silhouette method, where the optimal number of clusters is the one with the highest average silhouette width. Some publicly available genomes appeared to be misannotated according to their ANI-based clustering, specifically three *BL*. *longum* genomes clustered within the *BL. suis* group, and one *BL. infantis* genome was found within the *BL*. *longum* cluster. Previous reports have noted such mis-annotations, including for genome assembly GCA_000092325, which is likely misclassified as *BL. longum* and instead represents *BL. suis*^52,69^.

dDDH values were calculated using the genome-to-genome distance web-based calculator (GGDC 3.0^70,71^). Since full-genome dDDH analysis is computationally intensive, we selected a representative subset of 24 genomes, which included the type strains of all subspecies and all nine genomes belonging to the proposed *BL*. *nexti.* Genomes were uploaded to the web server, genomes were aligned using BLAST, distances and dDDH values were calculated using the d2 formula (identities / HSP length) of GGDC, which uses a generalized linear model (GLM) inferred from an empirical reference dataset comprising real DDH values and genome sequence^71^. To refine the phylogeny, core-gene alignment was performed in Roary^72^ (3.13.0) using the -e and --mafft flags. A maximum-likelihood tree was then constructed using RAxML^73^ (v8.2.12) with the GTRCAT model and the SSE3 protocol, employing the -f a flag for rapid bootstrap analysis.16s rRNA gene sequence was retrieved from genomes using Barrnap (0.9)^74^. Multiple sequence alignment and distance calculation of 16S rRNA gene was performed using Clustal Omega (1.2.4)^75^. 16s rRNA and core gene phylogenetic trees were created using RAxML with raxmlHPC-SSE3^73^.

Genome synteny was assessed using minimap2 (2-2.24)^60^ with -ax flag asm20, and visualised using the gggenomes (1.0.1) R package^76^. Gene clusters were searched using Integrative Genomics Viewer (IGV)^77^ and visualised using gggenomes^76^. Genes annotated as hypothetical proteins by Prokka^78^ were further analyzed using BLASTx^79^ against non-redundant protein sequences (nr) database and HHpred^80^ to infer potential functions. *BL. nexti*-specific gene sequences were mapped against the Uniref90 database^81,82^ using the uniref_annotator.py script^83^, which utilizes DIAMOND^84^ for alignment. Uniref90 annotations were then assigned a gene Gene Ontology (GO) annotations using UniProt.

Gene annotation for all 1,220 MAGs and genomes was performed using Prokka^78^ (v1.14.6), which uses BioPerl^85^, GNU Parallel^86^, BLAST+^79^ and Prodigal^87^. Pangenomic analysis was subsequently conducted with Roary^72^ (v3.13.0) using the GFF3 files generated by Prokka to define core and accessory genes and a gene presence/absence matrix.

To assess intra-subspecies diversity, we calculated Euclidean distances between genomes based on a gene presence/absence matrix. For each *B. longum* subspecies, we calculated the distance from the centroid for each group using the dist_to_centroids function from the usedist R package. To control for differences in sample size between subspecies, we performed random subsampling by selecting 34 genomes (corresponding to the size of the smallest subspecies group, *BL. suis*) from each subspecies. This procedure was repeated over 1,000 iterations, and results were averaged across all iterations to obtain robust estimates of centroid distances.

Downstream analyses were performed in R (v4.4.0) using tidyverse^88^, including dplyr^89^ (1.1.4), tidyr^90^ (1.3.2), stringr^91^ (1.5.1) and tidyverse^88^ (2.0.0). Genomic distances were determined by calculating euclidean distances from the Roary gene presence/absence matrix using the dist function from the stats package. Isolates were clustered via hierarchical clustering using the Ward.D2 method, and Principal Component Analysis (PCA) was performed using the ape (v5.8-1) package. Clade boundaries within the PCA space were defined using k-means clustering (k=4). Visualizations were generated using ggplot2^92^ (4.0.1), ggforce^93^ (0.5.0), and RColorBrewer^94^ (1.1-3). Heatmaps were produced with pheatmap^95^ (1.0.13), using the cutree function to divide into clusters. Phylogenetic trees were visualized and annotated using ggtree^96^ (3.12.0). Dendrograms were constructed using a neighbour joining algorithm with the nj function from the ape package^97^ (5.8.1) and visualized using ggtree^96^ (3.12.0). t-tests were performed using R.

### Microscopy

Culturing and microscopy was performed on representative strains of *BL. longum*, *BL. infantis* and three *BL. nexti* isolates; MB0401, MB0413 and LL6991^T^. *BL. infantis* and *BL. longum* were obtained from the BMB cohort^30^. Strains were grown routinely at 37°C in an anaerobic chamber (COY) that was maintained at > 2.5% H2 by flushing with a gas mix containing 20% H2, 5% CO2, and 75% N2. Strains were grown in supplemented Brain Heart Infusion (sBHI) media, supplemented with 50 ml/L fetal bovine serum (FBS; Sigma, F2442), 10 ml/L trace vitamins (ATCC® MD-VS™), 10 ml/L trace minerals (ATCC® MD-TMS™), 10 ml/L vitamin K1 and hemin (BBL, 212354), 1 g/L D-(+)-Cellobiose (Alfa Aesar, 528507), 1 g/L D-(+)-Maltose (Caisson, 6363537), 1 g/L D-(+)-Fructose (Sigma Aldrich, 1286504), and 0.5 g/L L-Cysteine (Acros Organics, 52904). For imaging, strains were cultured overnight and then diluted 30-fold in fresh medium and grown to exponential phase. Next, they were stained with DAPI (4-,6-diamidino-2-phenylindole, Sigma-Aldrich Cat #D8417) and FM 4-64 Dye (N-(3-Triethylammoniumpropyl)-4-(6-(4-(Diethylamino) Phenyl) Hexatrienyl) Pyridinium Dibromide, Invitrogen Cat# T13320). Cells were placed on an agarose pad with uncoated coverslips and imaged using a Nikon Eclipse Ti-E inverted microscope equipped with Perfect Focus System (PFS) and ORCA Flash 4 camera (Hamamatsu photonics). Images were processed, two-dimensional (2D) deconvolution and bacteria cell length measurement was performed using NIS Elements-AR software.

### DNA extraction and sequencing of vaginal samples

For vaginal swabs DNA was isolated from using the DNeasy PowerSoil Pro Kit (47016, Qiagen, Venlo, Netherlands), as described previously^98^. Vaginal swab samples were thawed at 4°C for 1 hour. Each sample was then vortexed for 3 minutes in 500 µL of PowerBead Solution. The liquid was transferred to PowerBead Pro Tubes, the total volume adjusted to 800 µL using CD1 solution and then vortexed briefly. Prepared vaginal samples were incubated at 65°C for 10 minutes and subsequently bead-beat at 5,000 rpm at 4°C for 45 sec on a Precellys Evolution tissue homogenizer (Bertin Instruments, Rockville, Maryland, USA). Samples were then centrifuged at 15,000 x *g* for 1 min at 4°C, and a 600 µl sample was used for automatic DNA extraction on Qiacubes (Qiagen) using the ‘DNeasy PowerSoil Pro Kit with Inhibitor Removal Technology Protocol’. DNA was eluted in 50 µl and stored at -20°C. Metagenomic sequencing vaginal microbiome was performed using the same protocol as for the gut samples as described previously^57^ with the only difference being that that the vaginal samples were sequenced at a higher depth (40 GB per sample) to account for the high proportion of human reads in this niche. Profiling was carried out using the same protocol as gut samples^57^.

### Subspecies-specific marker gene selection

To identify subspecies-specific genomic markers, we utilized the gene presence/absence matrix generated by Roary^72^. Gene sequences were functionally annotated by mapping against the UniRef90 database^81,82^ using the uniref_annotator.py script^83^, which employs DIAMOND^84^ for high-throughput protein alignment. Subspecies-specific markers were defined for *BL. infantis*, *BL. longum*, and *BL. nexti* based on a dual-threshold approach: genes were required to be present in >90% of genomes within the target subspecies and in <1% of genomes from each of the other subspecies. Given the high genomic similarity between *BL. longum* and *BL. suis*, marker genes for *BL. longum* were calculated without excluding sequences shared with *BL. suis*. As an extra reassurance that the analyzed genomes belonged to *B. longum*, we mapped MetaPhlAn 4.0 *B. longum* species-level marker genes and confirmed that >90% of genomes contained each of these markers, reinforcing the correct species assignment (**Figure 2C**, black gene annotation).

To quantify *B. longum* subspecies abundance in metagenomic datasets, we integrated these identified marker genes into the MetaPhlAn marker gene database (June 2023 release)^29^. This approach enables accurate high-resolution quantification of *B. longum* subspecies, as we previously showed^30^. We used this method to estimate the relative abundance of *BL. infantis*, *BL. nexti*, and *BL. longum* (the latter partially encompassing *BL. suis* and other possible unclassified members of the clade). In addition,while our classification method effectively distinguished subspecies-level reads, a portion of *B. longum* reads could not be assigned to a specific subspecies with high confidence and were categorized as unclassified *B. longum*. Since these unclassified reads exhibited longitudinal trends and distribution patterns highly consistent with *BL. longum* (**Supplementary Figure 6A-C**), they were aggregated with *BL. longum* for subsequent analyses.

### α-amylopullulanase gene analysis

To identify α-amylopullulanase homologs, the gene from *BL. nexti* was used as a query to search against 11,141 *Bifidobacterium* genomes from StrainAtlas using BLAST. For genomes containing multiple hits, only the highest-scoring locus was retained, resulting in the identification of α-amylopullulanase in 80 genomes. A rooted species phylogeny was inferred for these 80 genomes using GTDB-Tk. For gene tree reconstruction, α-amylopullulanase nucleotide sequences were aligned with MAFFT (v7.520; --auto), and a maximum-likelihood gene tree was estimated using FastTree in nucleotide mode (-nt). To ensure the gene tree orientation was consistent with the species-tree context, optimal rooting was performed with OptRoot in Ranger-DTL using the combined input of the species and gene trees. For reference-based divergence profiling, the α-amylopullulanase sequence from genome LL6991 served as the reference for comparison against the remaining 79 aligned sequences. At each alignment position, only pairs where both the reference and the query contained unambiguous nucleotides (A/C/G/T) were considered comparable; gaps or ambiguous bases (e.g., N) were treated as missing data and excluded from the analysis. Per-site divergence was calculated as the proportion of comparable genomes differing from the reference at a given position, with values averaged within non-overlapping 10-bp windows for visualization. Finally, genome synteny between *B. breve* UCC2003 and *BL. nexti* LL6991 was assessed using minimap2 (v2.24) with the -ax asm20 flag and visualized using the gggenomes (v1.0.1) R package.

Ancestral state reconstruction was assessed using gene presence/absence readouts (blastn hits were required to have e-value ≤ 0.01; among genomes with a hit, query coverage ≥40% was considered as present, genomes with no hit passing the e-value threshold were considered absence, taxa with intermediate results were removed) and a GTDB-Tk maximum likelihood phylogenetic tree of 11,151 StrainAtlas bifidobacteria, including 7 *BL. nexti* representatives, rooted in the outgroup lineage *Minisyncoccus archaeiphilus*. The R package castor v1.8.5 was used for maximum parsimony ancestral state reconstruction.

The α-amylopullulanase protein sequence was analyzed for functional domains and localization signals. Signal peptides were predicted using SignalP^99^ (6.0), and potential cell-wall anchoring motifs were identified through manual inspection of the sequence.

To evaluate the expression of α-amylopullulanase in starch, RT-qPCR was performed. *BL. nexti* DMP strain was cultured in YCFA supplemented with 0.05% cystine in two separate carbon sources namely 20 mM glucose and 20 mM water-soluble Difco starch (Catalog ID 217820; Becton Dickinson and Company, MD, USA). The cultures were grown in triplicates till mid-exponential phase (0.6-0.8) OD_600_. The harvested cultures were centrifuged at 4,000xg for 10 min at 4 °C . The supernatant was discarded and the pellets were stored in -80 °C after resuspending in 100 uL of Qiagen RNAprotect bacteria reagent (Qiagen,Hilden, Germany). On the day of RNA isolation, the cells suspended in the RNA protect agent were centrifuged at 4,000xg for 10 min and the supernatant was discarded. The RNA was then isolated using the Qiagen RNeasy mini kit (Qiagen,Hilden, Germany) with an additional bead beating step. 600 µL of RLT lysis buffer was mixed with the cell pellet and transferred to a 2ml screw cap microtube containing 0.1mm Zirconia/silica beads (BioSpec, Carl Roth). The bead beating step was conducted using Precellys Evolution Homogenizer with a setting of 6800 rpm for 3 cycles of bead beating for 20s and a pause of 30s in between each cycle. These tubes were then centrifuged at 14,000xg for 10 mins and the supernatant was transferred to a 2 mL microtube where 600 µL of 70% ethanol was added and mixed well by pipetting. A volume of 700 µL was then transferred to a clean 2 mL collection tube and the RNA isolation protocol was continued according to the manufacturer’s protocol. The RNA was eluted using 30 µL of RNase - free water and RNA quantity was measured.

500 ng of RNA was treated to remove DNA contamination using AMPD1 DNase kit (AMPD1-1KT,Sigma Aldrich,Massachusetts,USA) according to manufacturer’s protocol. The cDNA was then generated using reverse transcription from the DNA-free RNA as template using RevertAid First Strand cDNA Synthesis Kit (K1622, Thermo Scientific, Netherlands). The cDNA quantity was then diluted to 5ng/uL and RT-qPCR were carried out using iTAQ universal SYBR green supermix (Catalog ID 1725121,Bio-Rad Laboratories, Hercules, CA, USA). All qPCR reactions were run in triplicates using a 384-well plate on a QuantStudio™ 7 Flex (Thermo Scientific, Massachusetts, USA) with manufacturer’s thermo-cycler conditions. The primers designed for target α-amylopullulanase (FW-5’CAAGTTCCTTGGCATGGTGC3’,RW-5’GTAGTCGACGCCAGTAGACG3’, product size-76 bp) with housekeeping gene DNA gyrase(gyrA) (FW-5’CGTACCAAGATCCTGCCGTT3’, RW-5’TGACCACCACGTTCTCTTCG3’, product size-73 bp) to correct for initial RNA quantity variability. 2^−ΔΔCT^ method was used to calculate the relative change in expression of α-amylopullulanase between starch versus glucose.

### Carbohydrate utilization gene profiling and functional assays

To characterize the metabolic potential of the *B. longum* subspecies, we performed comprehensive carbohydrate utilization gene profiling across all genomes. We identified carbohydrate-active enzymes (CAZymes) using dbCAN3^100–102^. Specifically, we analyzed the results from the DIAMOND search against the CAZy database. To further investigate the evolutionary context of the identified α-amylopullulanase in *BL*. *nexti*, we conducted comparative protein sequence analyses. We used BLASTP^79^ to search for homologous proteins across other *Bifidobacterium* species.

Functional carbohydrate utilisation profiles of *BL. nexti* strains were compared to other *B. longum* subspecies. The *BL. nexti* strainsLL6991 (isolated from LLNEXT cohort), MB0401 (isolated from MicrobeMom cohort) and NextiDMP (Dutch microbiome project adult cohort) were used for strain level comparison. Type strains *BL. longum* E194b and *BL. infantis* S12 were used for subspecies comparison. To test the specific growth of these strains on various carbohydrates they were grown on YCFA supplemented with 20 mM autoclaved (120 °C for 20 mins) water-soluble Difco starch (Catalog ID 217820; Becton Dickinson and Company, MD, USA), pullulan (Catalog ID - P4516, Darmstadt, Germany) along with 20 mM glucose as positive and negative control. The cultures were incubated for 48 hours at 37°C in triplicate (Becton Dickinson and Company, MD, USA) and OD600 readings taken every 30 mins using Biotek Epoch2 microplate reader. All analyses were performed in R (version 4.4.1) using tidyverse, readr, ggplot2, scales,viridis, ggdendro and cowplot packages.

### Global Metagenomic Dataset Curation and Profiling

To evaluate the global prevalence and abundance of *B. longum* subspecies across diverse populations, we compiled public fecal metagenomic data spanning 18 distinct geographic locations (6,162 samples). Public cohorts were selectively included based on the availability of high-quality shotgun metagenomic sequencing data from both infant and adult populations, representing varying clinical and dietary backgrounds. Human contamination was filtered by mapping reads against the human reference genome (hg38) using Bowtie2 (2.4.5-1). To accurately distinguish and quantify individual *B. longum* subspecies, including the newly proposed *BL. nexti*, we profiled the clean metagenomic reads using our customized MetaPhlAn database augmented with the unique subspecies-specific marker genes identified in this study. This custom database was applied to infant stool metagenomes from geographically diverse cohorts, including Bangladesh^19^, Israel^30^, the USA (representing datasets from Boston^103^, Pennsylvania^104,105^, California^104^, Georgia^104^, Oregon^104^, and South Carolina^104^), Estonia^106^, Finland^106^, Russia^106^, Ireland^55^, Luxembourg^107^, the Netherlands^31^, the United Kingdom^54^, Spain^108^, and Sweden^7^.

### Assessment mother-infant dominant strain similarity

We used the MetaPhlAn alignments to marker genes to extract dominant-strain haplotypes using MetaPhlAn’s sample2marker script. For those families where we identified taxonomic presence in either *BL. infantis* or *BL. nexti* for at least one time point in both infant and mother, we extracted their respective marker genes, and ran a multiple sequence alignment per marker gene using MAFFT v7.520. Then, per subspecies, we computed overall Hamming distances, and the percentage of identity between samples. We filtered out pair comparisons that did not reach at least 800 nucleotides.

To test whether maternal samples were more similar to their infants than unrelated samples, we annotated samples as mother-infant dyads or unrelated (no dyad and no longitudinal). Then, we tested the hypothesis that the distribution of similarity between both groups were different, by using a Wilcoxon signed-rank test, approximating the null distribution throughout 10,000 permutations, as implemented in R’s coin v1.4.3 package. In order to account for the hierarchical nature of the data (multiple comparisons of the same subjects at different timepoints), we kept only the sample comparison with the highest similarity per subject pair. Only 6 mother infant dyads could be tested for *BL. nexti*, while none were available for *BL. infantis*.

### Association analysis

We performed associations between host phenotypic data and *B. longum* subspecies using the Dutch LLNEXT cohort. Taxa relative abundances were first center-log-ratio (CLR) transformed. For each subspecies, we applied two distinct statistical approaches per phenotype (**Supplementary Table 7**). Model 1 tested the effect of the phenotype of interest in the CLR-transformed abundance of the subspecies using a linear mixed model with a random intercept per infant. Model 2 utilized logistic regression to test the effect of phenotypes on the "ever presence" of a subspecies (abundance > 0 at any timepoint). Model 2 was not applied to *BL. longum*, as nearly all infants were colonized at some point during their first year. To evaluate maternal transmission, we identified mothers with the presence of each subspecies during pregnancy (abundance > 0) and applied Model 1 to early infant timepoints (W2, M1, M2, or M3). Temporal abundance changes were assessed using Model 1 across the full one-year time series.

Delivery mode was tested using both Models 1 (accounting for time) and 2. Delivery place was tested only within vaginally delivered infants while accounting for time, as all cesarean deliveries occurred in hospital settings. Parity was binarized (no siblings vs. one or more siblings) and tested using Model 1, accounting for time and delivery mode. For feeding mode in *BL. nexti*, we expanded Model 1 to include an interaction term between time and delivery mode; we then performed a likelihood ratio test against the basic Model 1 to assess if the interaction term significantly improved model fit. For feeding mode associations in Model 2, we included only exclusively breastfed or exclusively formula-fed infants. Associations for *BL. infantis* were analyzed across the full year, whereas *BL. nexti* and *BL. longum* analyses were restricted to early timepoints (W2 to M3).

We additionally tested whether the presence of *BL. nexti* in maternal vaginal samples (presence/absence) increased the odds of infant’s ever presence, using a logistic regression (Model 2) as previously explained.

## Acknowledgements

The data used in this manuscript are provided by LLNEXT. The authors are grateful for the participation of all the parents and infants in LLNEXT. We thank the LLNEXT team and maternity care providers for recruiting participants, collecting material during and after childbirth and building and maintaining the cohort. We also thank the Genomics Coordination Center and the Center for Information Technology of the University of Groningen for their support and for providing access to the Gearshift, Peregrine, and Habrok high-performance computing clusters.

The Lifelines NEXT cohort study received funds from the University Medical Center Groningen Hereditary Metabolic Diseases Fund, Health∼Holland (Top Sector Life Sciences and Health), the EU, the Northern Netherlands Alliance (SNN), the provinces of Friesland and Groningen, the municipality of Groningen, Philips, and the Société des Produits Nestlé.

A. Z. is supported by the Nederlandse Organisatie voor Wetenschappelijk Onderzoek-VIDI (NWO-VIDI) grant 016.178.056, and NWO-VICI grant VI.C.232.074, EU Horizon Europe Program grant INITIALISE (101094099), NWO KIC grant KICH1.LWV04.21.01and NWO Gravitation grant Exposome-NL (024.004.017). M.Y. is the Rosalind, Paul and Robin Berlin Faculty Development Chair in Perinatal Research and is supported by the Azrieli Foundation grant for faculty fellows. J.F. is supported by the ERC Consolidator grant (grant agreement No. 101001678), NWO-VICI grant VI.C.202.022, the AMMODO Science Award 2023 for Biomedical Sciences from Stichting Ammodo.

---

**Supplementary Figure 1: *Bifidobacterium longum* comprises four distinct clades. (A)** Average genomic ANI (average nucleotide identity) between all genomes and MAGs. Genomes are colored based on prior public subspecies annotations, while MAGs from the LLNEXT cohort are colored according to their sample annotation as defined in the phylogenetic tree in Figure 1A. **(B)** Heatmap showing digital DNA-DNA hybridization (dDDH) values among 24 selected genomes. **(C)** Phylogenetic tree of 24 selected genomes based on the 16S rRNA gene nucleotide sequences, with *B. breve* used as the outgroup. Subspecies clustering is not resolved because the 16S rRNA gene lacks sufficient sequence variation to distinguish between *B. longum* subspecies. **(D)** Microscopic images oftwo additional *BL. nexti* strains during the logarithmic growth phase. Cells are stained to visualize the cell membrane and DNA. Scale bar, 2.5 μm.

**Supplementary Figure 2: Gene content profiles distinguish the four *Bifidobacterium longum* subspecies.** Presence of pangenome genes across all genomes and MAGs. The dataset was filtered to include only genes present in at least five genomes or MAGs. Genomes and MAGs are annotated based on prior public annotations or the marker-gene-based phylogenetic tree from Figure 1A.

**Supplementary Figure 3: Genomic diversity and synteny among *Bifidobacterium longum* subspecies. (A)** Within-subspecies genomic variability relative to the corresponding type strain. Variability was calculated as the Euclidean distance between each genome or metagenome-assembled genome (MAG) and the type strain of its assigned subspecies based on gene presence/absence profiles. Genomes and MAGs were assigned to subspecies using pangenome-based clustering. ****, *P* ≤ 0.0001. **(B)** Genomic synteny among the type strains of the four *Bifidobacterium longum* subspecies. Each horizontal line represents a genome, with multiple lines indicating separate contigs. Green shading denotes conserved collinear regions identified by whole-genome sequence alignment. *BL. nexti*-specific genes are highlighted in orange.

**Supplementary Figure 4: Comparative sequence divergence and genomic synteny of α-amylopullulanase. (A)** Species-stratified sequence divergence from the *BL. nexti* LL6991 α-amylopullulanase reference. For each position in the multiple sequence alignment, divergence was calculated relative to the LL6991 reference as the fraction of comparable genomes (those with unambiguous A/C/G/T in both reference and query) differing from the reference. Sites containing gaps or ambiguous bases were excluded. Divergence values were averaged within 10-bp sliding windows and plotted per species. Windows lacking comparable sites within a species are omitted. The y-axis is fixed from 0 to 1, where 0 indicates complete identity and 1 indicates that all comparable genomes differ from the reference. **(B)** Genomic synteny of the α-amylopullulanase locus , comparing the genomic region surrounding the α-amylopullulanase gene in the *BL. nexti* type strain and *B. breve* UCC2003. The green shaded area highlights an 11-kb region of high similarity between the two genomes. Homologous genes are indicated by matching colors. **(C)** Ancestral state reconstruction of the α-amylopullulanase gene distribution. Maximum-likelihood GTDB-Tk phylogeny of *Bifidobacterium* genomes showing the subset containing *B. breve*, *B. longum*, and *BL. nexti*. Tips are colored according to species, and shapes indicate the presence or absence of the α-amylopullulanase gene. Ancestral gene states were inferred using maximum parsimony reconstruction based on gene presence/absence calls across the full phylogeny.

**Supplementary Figure 5: Growth dynamics and α-amylopullulanase expression in *BL. nexti*. (A)** Growth curves (OD_6 00_) of seven strains representing *B. longum* subspecies and *B. breve* over 48 hours on glucose (positive control), pullulan, starch, and 2′-fucosyllactose (2′FL). Curves represent the mean of three biological replicates, with substrate-specific negative controls included. **(B)** Growth curve (OD_600_) of *BL. nexti* (strain LL6991) grown on different carbohydrates, highlighting its ability to utilize glycogen. **(C)** Relative expression of the α-amylopullulanase gene in *BL. nexti* (strain DMP118) during growth on starch compared with glucose, measured by RT-qPCR and normalized to a housekeeping gene. Gene expression is shown as fold change relative to growth on glucose. Points represent three independent biological replicates (each measured in technical triplicate)

**Supplementary Figure 6: Longitudinal dynamics and co-occurrence patterns of *B. longum* subspecies. (A-B)** Longitudinal relative abundance of *B. longum* subspecies in the LLNEXT cohort. Boxplots represent the relative abundance of *BL. longum* (including unclassified *B. longum*) in **(A)** mothers and **(B)** infants across multiple time points. **(C)** Scatter plot showing the relative abundance of *BL. longum* versus unclassified *B. longum* across all metagenomic samples. The strong linear correlation suggests that unclassified *B. longum* largely represents *BL. longum* lineage that is not captured by our classification markers. **(D-E)** Subspecies co-occurrence patterns. Scatter plots illustrating the relative abundance of *BL. nexti* versus **(D)** *BL. infantis* and **(E)** *BL. longum* across all infant samples. The distributions demonstrate a pattern of mutual exclusivity between *BL. nexti* and other *B. longum* subspecies.

**Supplementary Figure 7: Association of early-life factors with the presence of *B. longum* subspecies in the infant gut microbiome. (A-C)** Longitudinal prevalence of *B. longum* subspecies stratified by birth environment and parity. Plots illustrate the prevalence over time for **(A-B)** *BL. longum* and **(C)** *BL. nexti*. Infants are stratified by **(A)** birth setting among vaginally delivered infants (home vs. hospital delivery) and **(B, C)** maternal parity (firstborn infants [0] vs. non-firstborn infants [≥1]). Absolute numbers for each group (N) are indicated at the top of each plot.

**Supplementary Figure 8: Subspecies co-occurrence patterns.** Scatter plots illustrating the relative abundance of *B. adolescentis* versus *BL. nexti*, *BL. infantis* and *BL. longum* across all infant samples. The distributions demonstrate a pattern of mutual exclusivity between *B. adolescentis* and *BL. nexti*.

**Supplementary Table 1:** Summary of *Bifidobacterium longum* genomes and metagenome-assembled genomes (MAGs) together with metadata analyzed in this study.

**Supplementary Table 2: Pairwise ANI matrix.** Average Nucleotide Identity (ANI) values calculated for all *B. longum* genomes and MAGs used in the study.

**Supplementary Table 3: Metadata for *BL. nexti* isolates**. This table summarizes key metadata for the nine *BL. nexti* genomes analyzed in this study, including isolation source of the strains, geographic origin, and relevant genomic features.

**Supplementary table 4: Pairwise dDDH values for representative genomes.** Digital DNA-DNA hybridization (dDDH) matrix for the 24 representative genomes, including the type strains of all recognized *B. longum* subspecies and the genomes belonging to *BL*. *nexti*.

**Supplementary Table 5:** Catalog of subspecies-specific genes for *BL. longum*, *BL. infantis*, and *BL. nexti*.

**Supplementary Table 6:** List of 80 genomes from five distinct *Bifidobacterium* species found to encode α-amylopullulanase. The table includes species designations and the percent identity of each homolog relative to the α-amylopullulanase gene identified in the *BL. nexti* LL6991 representative strain.

**Supplementary Table 7:** Associations between *B. longum* subspecies and early-life phenotypes. Summary of statistical analyses examining the prevalence and relative abundance of *B. longum* subspecies in relation to infant metadata.

