## Supplementary figures and images for "Discovery, functionality and global dynamics of *Bifidobacterium longum* subsp. *nexti*: a novel starch-degrading subspecies in the infant gut"

### Supplementary Figure 1

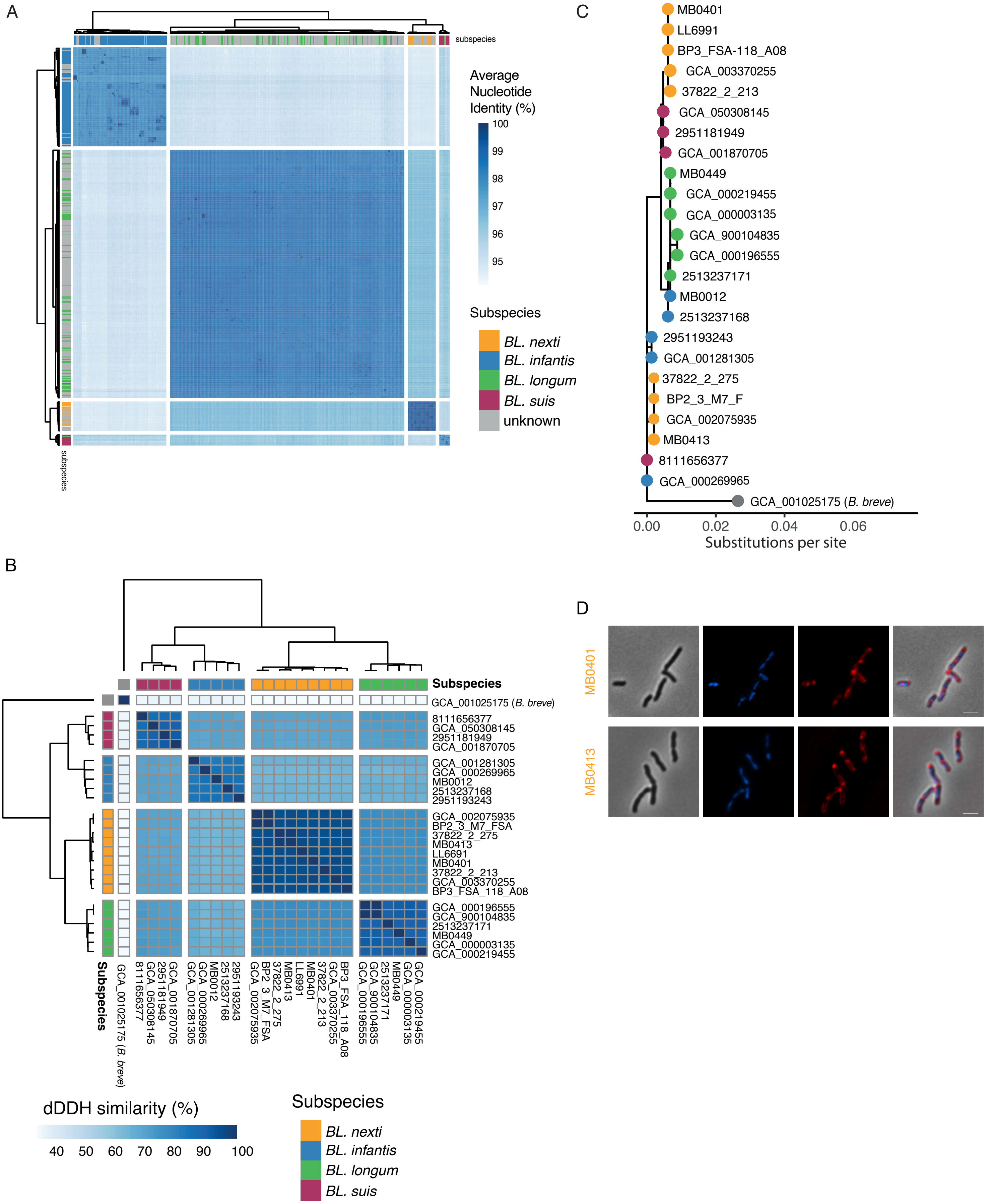

### Supplementary Figure 2

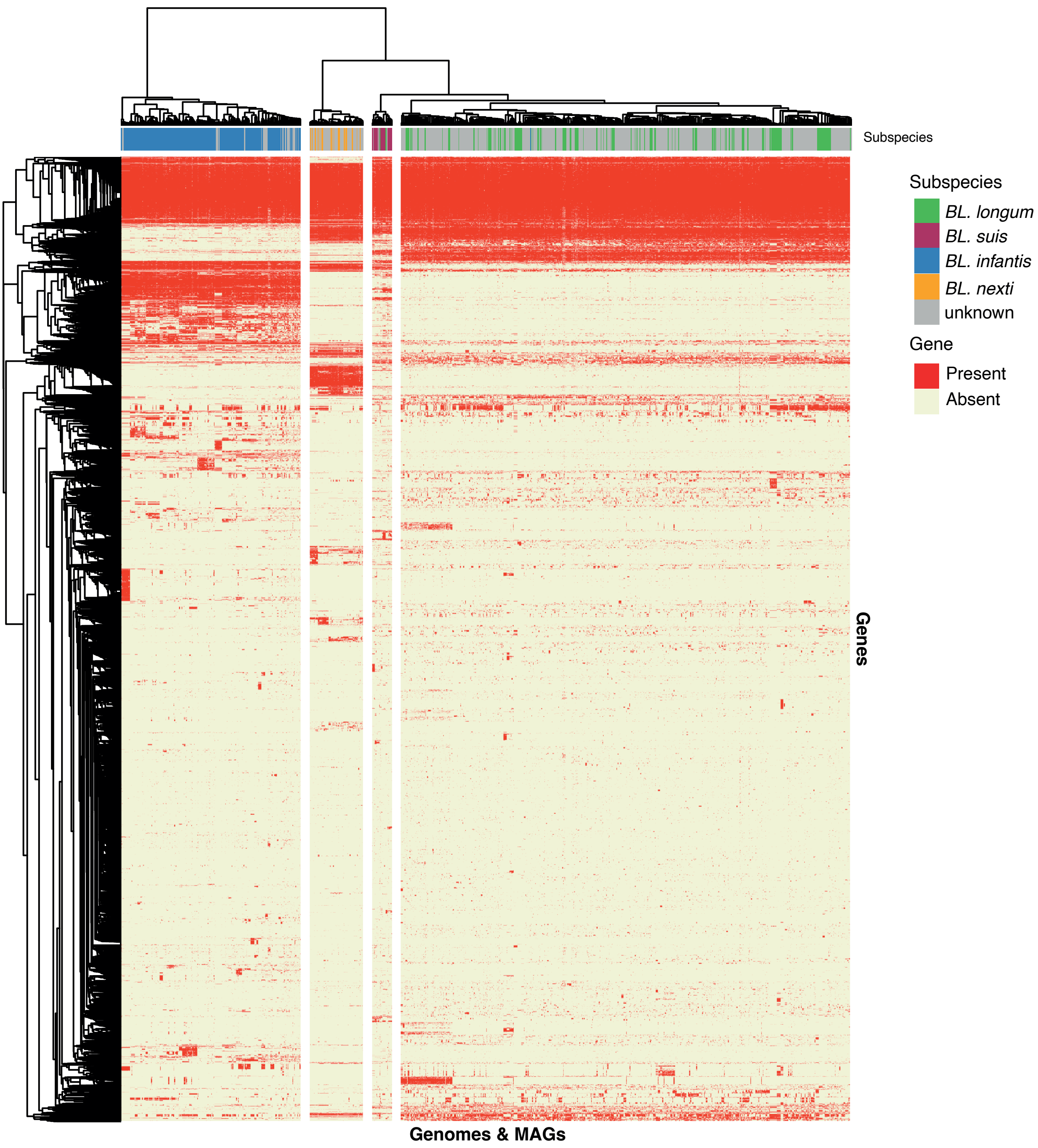

### Supplementary Figure 3

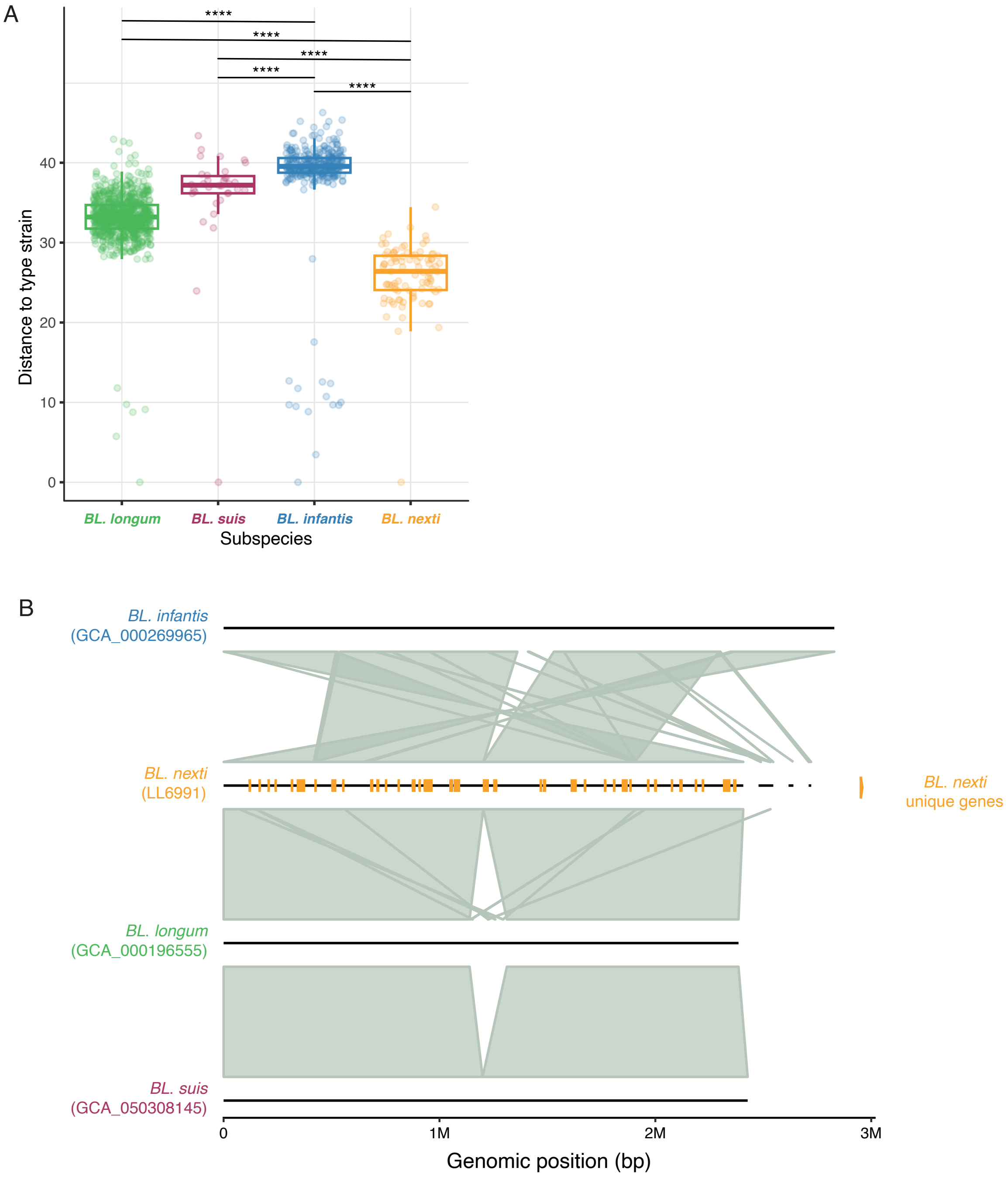

### Supplementary Figure 4

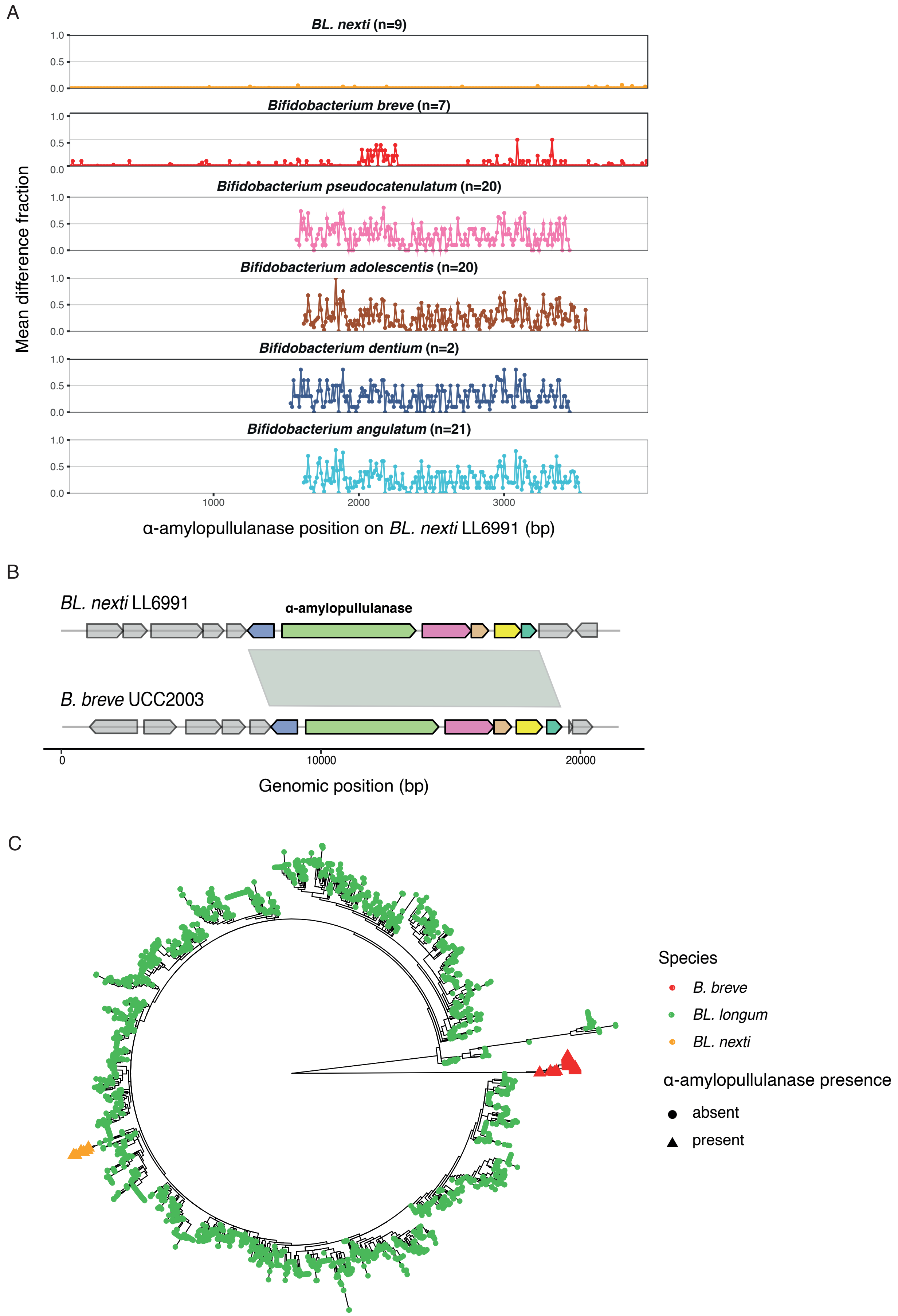

### Supplementary Figure 5

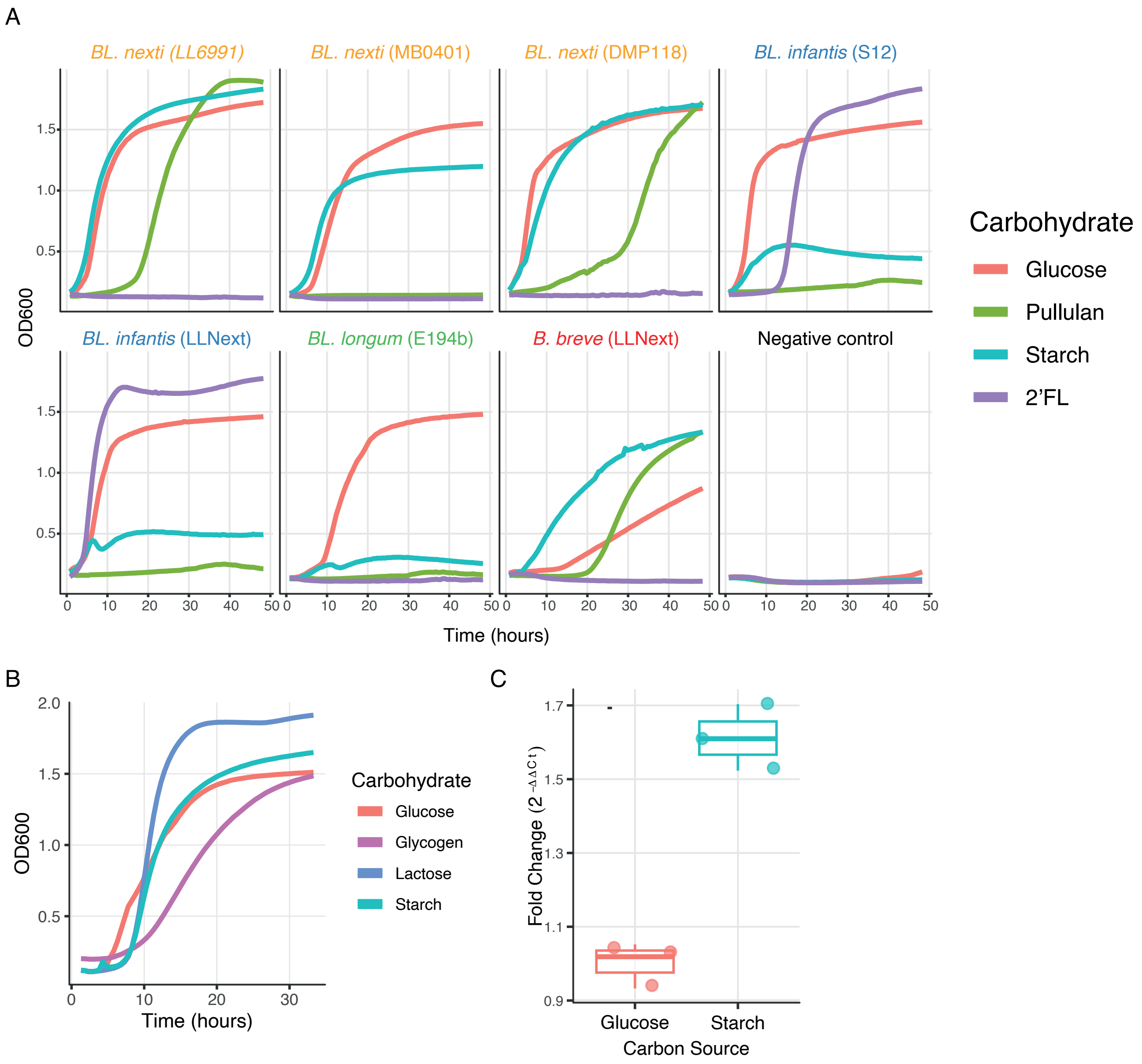

### Supplementary Figure 6

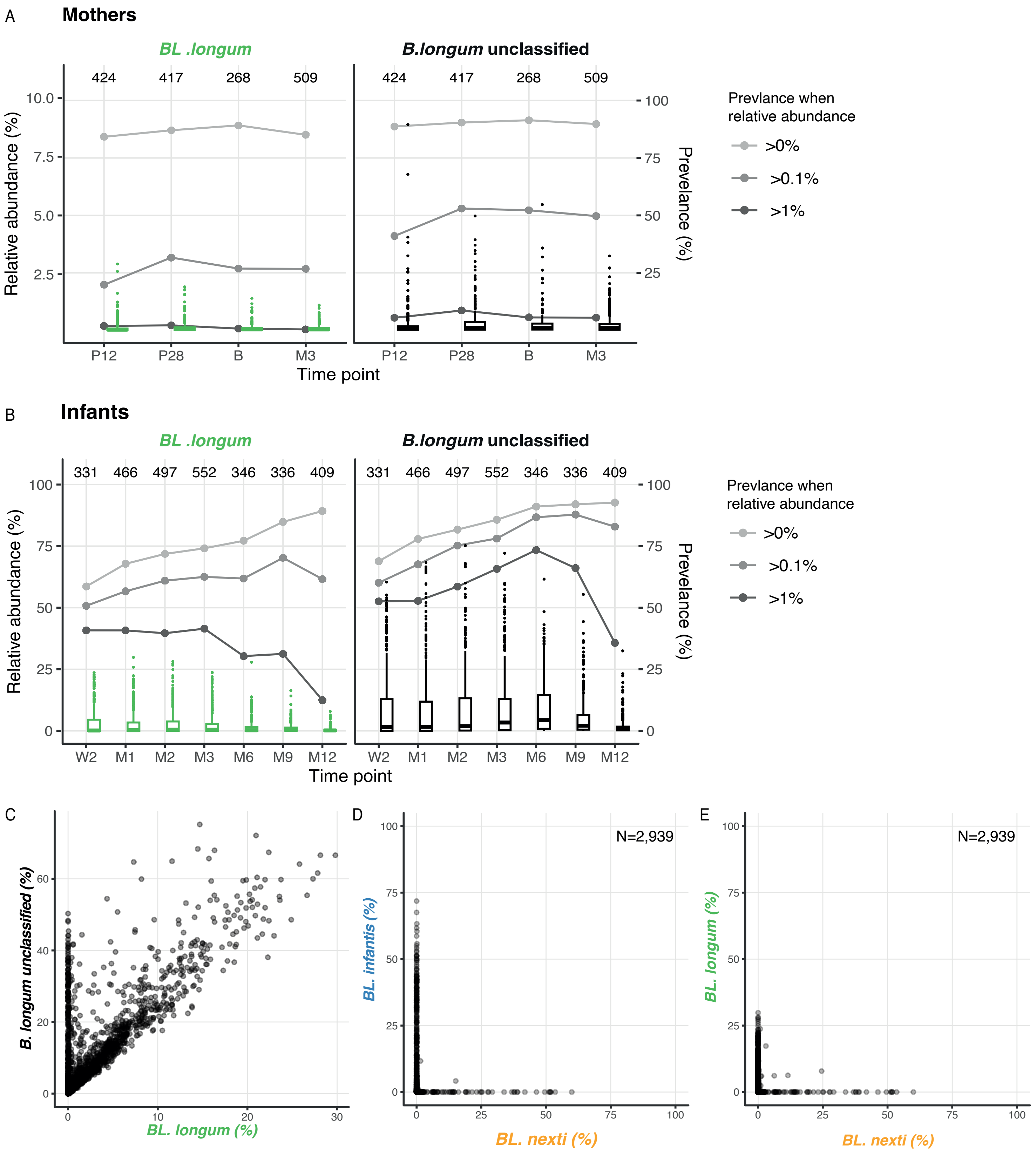

### Supplementary Figure 7

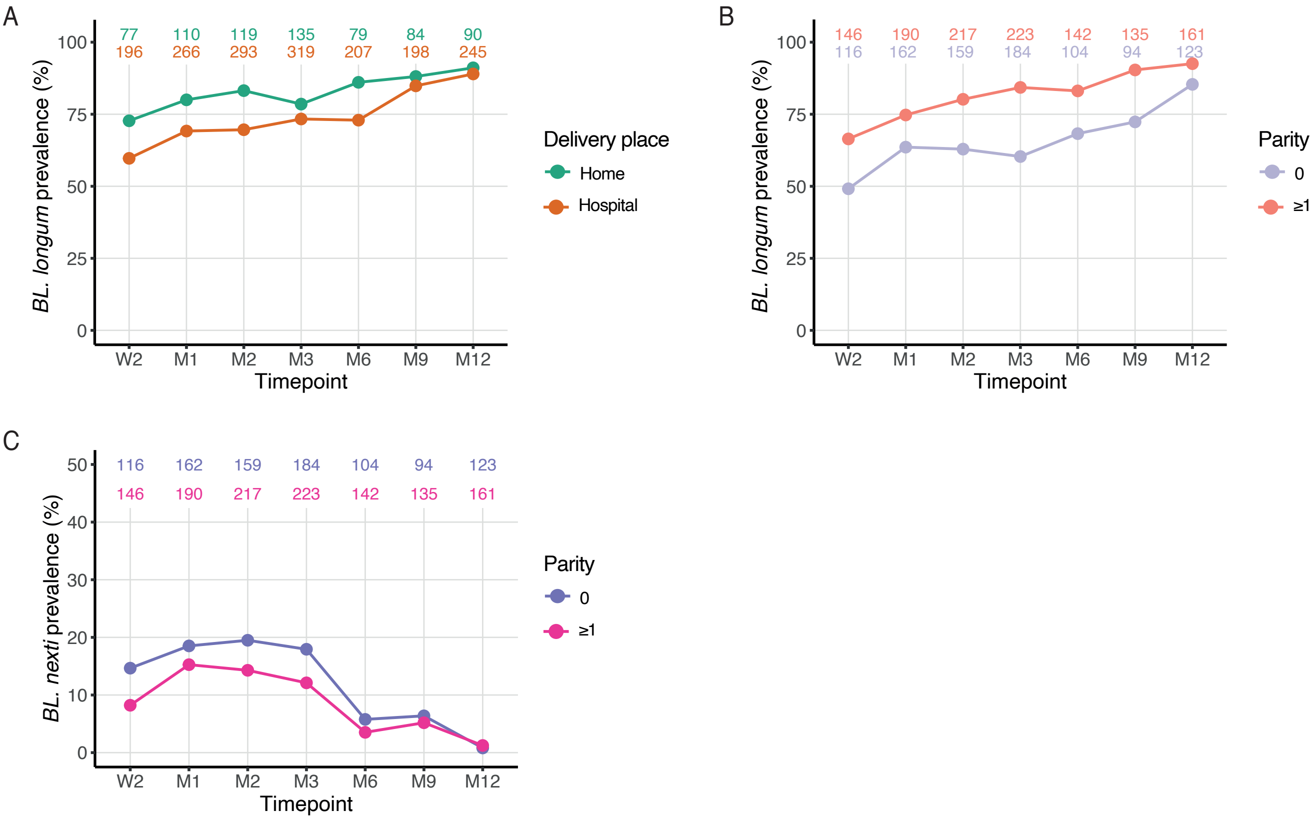

### Supplementary Figure 8

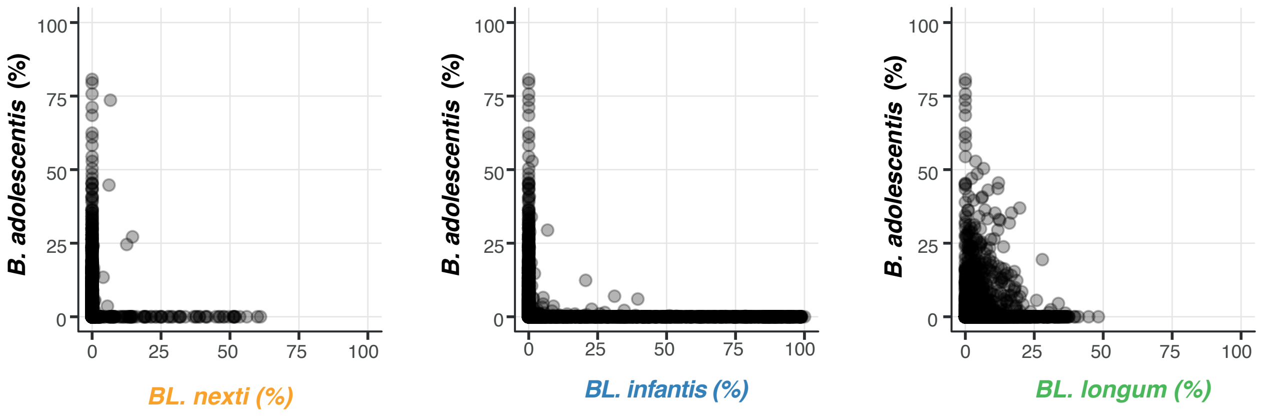
